# Structural insights into the assembly mechanism of the flagellar MS-ring with three different symmetries

**DOI:** 10.1101/2024.07.30.605930

**Authors:** Miki Kinoshita, Fumiaki Makino, Tomoko Miyata, Katsumi Imada, Keiichi Namba, Tohru Minamino

## Abstract

The flagellar basal body MS-ring, formed by 34 FliF subunits, is the core of the flagellar motor as well as the base for flagellar assembly. The MS-ring is also a housing for the flagellar protein export gate complex that is required for construction of the flagellum on the cell surface. A large periplasmic region of FliF contains three ring-building motifs named RBM1, RBM2, and RBM3. RBM3 forms the S-ring and β-collar with C34 symmetry. RBM2 forms the inner core ring of the M-ring with C23 symmetry surrounded by 11 cog-like structures formed by RBM1 and RBM2. However, it remains unknown how FliF assembles to generate these three different symmetries within the MS-ring. Here, we report the two cryoEM structures of the MS-ring formed by FliF co-expressed with FliG and transmembrane export gate proteins. Structural comparison of 33-mer and 34-mer MS-rings revealed that a subtle change in the conformation of RBM3 produces the different rotational symmetries. Combination of cryoEM structural and mutational analyses of the MS-ring with C33 symmetry provides evidence that the well-conserved DQxGxxL motif within a flexible loop connecting RBM2 and RBM3 allows RBM2 to take two different orientations relative to RBM3 to form not only 11 cog-like structures just outside the inner core ring with C22 symmetry but also an appropriately sized central pore in the inner core ring to accommodate the export gate complex.

**IMPORTANCE:** The flagellar MS-ring is the core of the flagellar motor and serves not only as an initial template for flagellar assembly but also as a base to accommodate the flagellar protein export complex. The MS-ring is formed by 34 subunits of FliF with two transmembrane helices and a large periplasmic region containing ring-building motifs, RBM1, RBM2, and RBM3. FliF adopts two different conformations in the MS-ring to generate three different rotational symmetries, C34, C23, and C11. However, how FliF assembles to produce these three symmetries remains a mystery. Combination of cryoEM structural and mutational analyses has provided evidence that the well-conserved DQxGxxL motif within a hinge loop connecting RBM2 and RBM3 allows RBM2 to take two different orientations relative to RBM3, allowing 23 RBM2 domains of 34 subunits to form the inner core ring with a properly sized central pore to accommodate the flagellar protein export complex.

## INTRODUCTION

The flagellum of *Salmonella enterica* serovar Typhimurium (hereafter referred to as *Salmonella*) is a supramolecular motility machine consisting of at least three structural parts: the basal body that acts as a bi-directional rotary motor; the filament that functions as a helical propeller; and the hook that works as a universal joint to smoothly transmit motor torque to the filament. The flagellar motor is placed under controls of intracellular chemotactic signaling networks, allowing bacterial cells to migrate towards environments suitable for survival (1,2).

The *Salmonella* basal body consists of the MS-ring (FliF), C-ring (FliG, FliM, FliN), LP-ring (FlgH, FlgI), and rod (FliE, FlgB, FlgC, FlgF, FlgG) (Fig. S1). These structures are highly conserved among bacterial species (3,4). The MS-ring is formed by 34 FliF subunits (5–7). FliG uses the MS-ring as a template for its assembly with 1:1 stoichiometry with FliF and forms the FliG ring on the cytoplasmic face of the MS-ring (8–11). Then, the FliM-FliN complex with a stoichiometry of 1 FliM subunit and 3 FliN subunits assembles onto the FliG ring through an interaction between FliG and FliM (12,13), forming the C-ring with C34 symmetry with a small symmetry variation (14–16). The MS-C-ring complex acts as a rotor of the flagellar motor. The MotA-MotB complex with a stoichiometry of 5 MotA subunits and 2 MotB subunits assembles around the rotor through an interaction between MotA and FliG and acts as a stator unit of the flagellar motor (17). The C-ring also functions as a directional switching device that allows the flagellar motor to rotate both in counterclockwise (CCW) and clockwise (CW) directions (18). The LP-ring with C26 symmetry functions as a molecular bushing for high-speed rotation of the rod acting as a drive shaft of the flagellar motor (5,6,19). The rod is a helical tubular structure with a helical symmetry of about 5.5 subunits per one turn of helix (20) and is divided into two structural parts: the proximal rod composed of 6 FliE, 5 FlgB, 6 FlgC, and 5 FlgF subunits; and a distal rod composed of 24 FlgG subunits (5,6,21,22). The proximal and distal rods are located inside the MS- and LP-rings, respectively. The basal body also contains the flagellar type III secretion system (hereafter referred to as fT3SS) that transports flagellar structural subunits from the cytoplasm to the distal end of the growing flagellar structure for construction of the flagellum on the cell surface (23). The fT3SS is composed of a transmembrane export gate complex with a subunit stoichiometry of 9 FlhA, 1 FlhB, 5 FliP, 4 FliQ, and 1 FliR and a cytoplasmic ATPase ring with a subunit stoichiometry of 12 FliH, 6 FliI, and 1 FliJ. The export gate complex is housed in the central pore of the MS-ring whereas the cytoplasmic ATPase complex is associated with the C-ring via FliH dimers as bridges (24,25). Thus, the MS-ring is the base for flagellar structure, assembly, and function.

*Salmonella* FliF (UniProt ID: P15928) consists of 560 amino acid residues with two predicted transmembrane helices (residues 26–46 and residues 455–475) and forms the M-ring, S-ring, and β-collar (26,27). The M-ring is embedded within the cytoplasm membrane whereas the S-ring and β-collar are in the periplasmic space. The C-terminal cytoplasmic domain of FliF is involved in the interaction with the N-terminal domain of FliG to connect the C-ring with the MS-ring (9). A large periplasmic region (residues 47–454) between the two transmembrane helices contains three ring-building motifs named RBM1 (residues 60–124), RBM2 (residues 125–215), and RBM3 (residues 228–438). RBM3 is further divided into three regions, RBM3a (residues 228–270), β-collar (residues 271–381), and RBM3b (residues 382–438). RBM3a and RBM3b form a single core domain (hereafter referred to as the S-ring domain) that assembles to form the S-ring with C34 symmetry. The β-collar is a cylindrical β-barrel structure above the S-ring. It consists of two sets of antiparallel β-sheets and firmly and stably accommodates the proximal rod. RBM2 is oriented in two distinct directions relative to the S-ring domain in the MS-ring. Of 34 FliF subunits, 23 RBM2 domains (RBM2_inner_) are placed inside the M-ring to form its inner core ring whereas the remaining 11 (RBM2_outer_) are located just outside the inner core ring together with RBM1 (RBM1_outer_). As a result, the export gate complex of the fT3SS is firmly and stably accommodated within the central pore of the inner core ring. Thus, FliF takes two distinct conformations in the MS-ring to generate three different rotational symmetries, C34, C23, and C11 (28,29). However, it is unclear how FliF dose so.

To address this question, we performed electron cryo-microscopy (cryoEM) structural analyses of the *Salmonella* MS-ring formed by FliF co-expressed with FliG and transmembrane export gate proteins and obtained two RBM3-ring structures with C33 and C34 symmetry at 2.4 Å and 2.5 Å resolution, respectively. We show that a conformational flexibility of the loops connecting the secondary structures within the S-ring and β-collar is responsible for slightly different angles between adjacent RBM3 domains to produce different rotationally symmetries without changing the interaction surfaces between domains. Iterative 3D refinement of the 33-mer MS-ring with C11 symmetry improved the resolution of the entire MS-ring structure from 4.2 Å to 3.1 Å. The structure reveals that conformational flexibility of a hinge loop connecting RBM2 and RBM3 (residues 214–228) allows FliF to take two different orientations of RBM2 relative to RBM3 so that 22 or 23 RBM2 domains form the inner core ring with a properly sized central pore to accommodate the flagellar protein export complex and the remaining 11 RBM2 domains form cog-like structures just outside the inner core ring, together with 11 RMB1 domains.

## RESULTS

### CryoEM structural analysis of the MS-ring formed by co-expression of FliF with FliG and transmembrane export gate proteins

The MS-ring is not only the structural template for FliG ring formation but also the housing for the transmembrane export gate complex made of FlhA, FlhB, FliP, FliQ, and FliR. FliO is required for efficient formation of the FliP_5_-FliQ_4_-FliR_1_ core structure before the export gate complex is incorporated into the central pore of the M-ring during MS-ring formation (30,31). Therefore, in this study, FliF was co-expressed with FlhA, FlhB, FliG, FliO, FliP, FliQ, and FliR from a pTrc99-based plasmid in *Salmonella* SJW1368 cells, in which the flagellar master operon named the *flhDC* operon required for the expression of all flagellar genes is missing, and fractions containing FliF, FliG, and FlhA were separated from solubilized crude membranes using sucrose gradient ultracentrifugation (Fig. S2A). Ring-shaped particles were observed in both negative stained EM images (Fig. S2B) and cryoEM images (Fig. S3A). In the flagellar basal body structure, FlhA is located inside the MS-ring along with other export gate proteins (30,32,33), and its C-terminal cytoplasmic domain forms a homo-nonameric ring structure that projects into the central cavity of the C ring (34,35). Although FliG and FlhA were associated with purified MS-ring (Fig.S2A), the FliG-ring and the export gate complex could not be observed by our cryoEM image analysis.

A total of 1,015,741 particles were extracted from 4,885 micrographs and analyzed. It has been shown that the MS-ring formed by co-expression of full-length FliF with FliG, FliM, and FliN has rotational symmetry ranging from 32-fold to 34-fold, but the 33-mer rings are the most commonly observed (about 80% of the population) compared to the 32-mer (about 5% of the population) and 34-mer rings (about 15% of the population) (36). In the present cryoEM analysis, representative 3D classification revealed that the ring structures fall into two distinct classes: 33-mer (70,250 particles, 53% of the population) and 34-mer (62,306 particles, 47% of the population) (Fig. S3B). No other symmetries were observed in the 3D class averages, indicating that the purified MS-ring consists of 33 or 34 FliF subunits. The MS-ring in the native basal body has 34-fold symmetry, and C-terminal truncations of FliF result in the symmetry variation in the MS-ring (5–7,28,36,37). The SDS-PAGE band pattern of our FliF sample indicates that it is full-length FliF that forms these MS-ring structures (Fig. S2A). Because FliF was overexpressed from the plasmid along with FliG, FlhA, FlhB, FliO, FliP, FliQ, and FliR, a small variation seen in the ring stoichiometry would be due to a proper or improper incorporation of the transmembrane export gate complex into the MS-ring.

We carried out 3D image reconstruction of the RBM3-ring containing 33 or 34 FliF subunits without rotational symmetry, and the resolution reached 4.2 Å and 4.6 Å for the 33-mer and 34-mer rings, respectively (Fig. S3B). We then performed iterative 3D refinements with C33 and C34 symmetry, respectively. This process dramatically improved the resolution of the RBM3-ring structure to 2.4 Å and 2.5 Å for the 33-mer (EMDB ID: EMD-60008) and 34-mer (EMDB ID: EMD-60009), respectively (Fig. 1A and Fig. S3C), allowing us to construct accurate atomic models of the 33-mer (PDB ID: 8ZDT) and 34-mer (PDB ID: 8ZDU) (Fig. 1B). These two atomic models were nearly identical to the corresponding parts of the previously reported *Salmonella* MS-ring structures (Fig. S4), but the map resolutions were slightly better than the previous ones (5–7,27,35,36).

**FIG 1.**
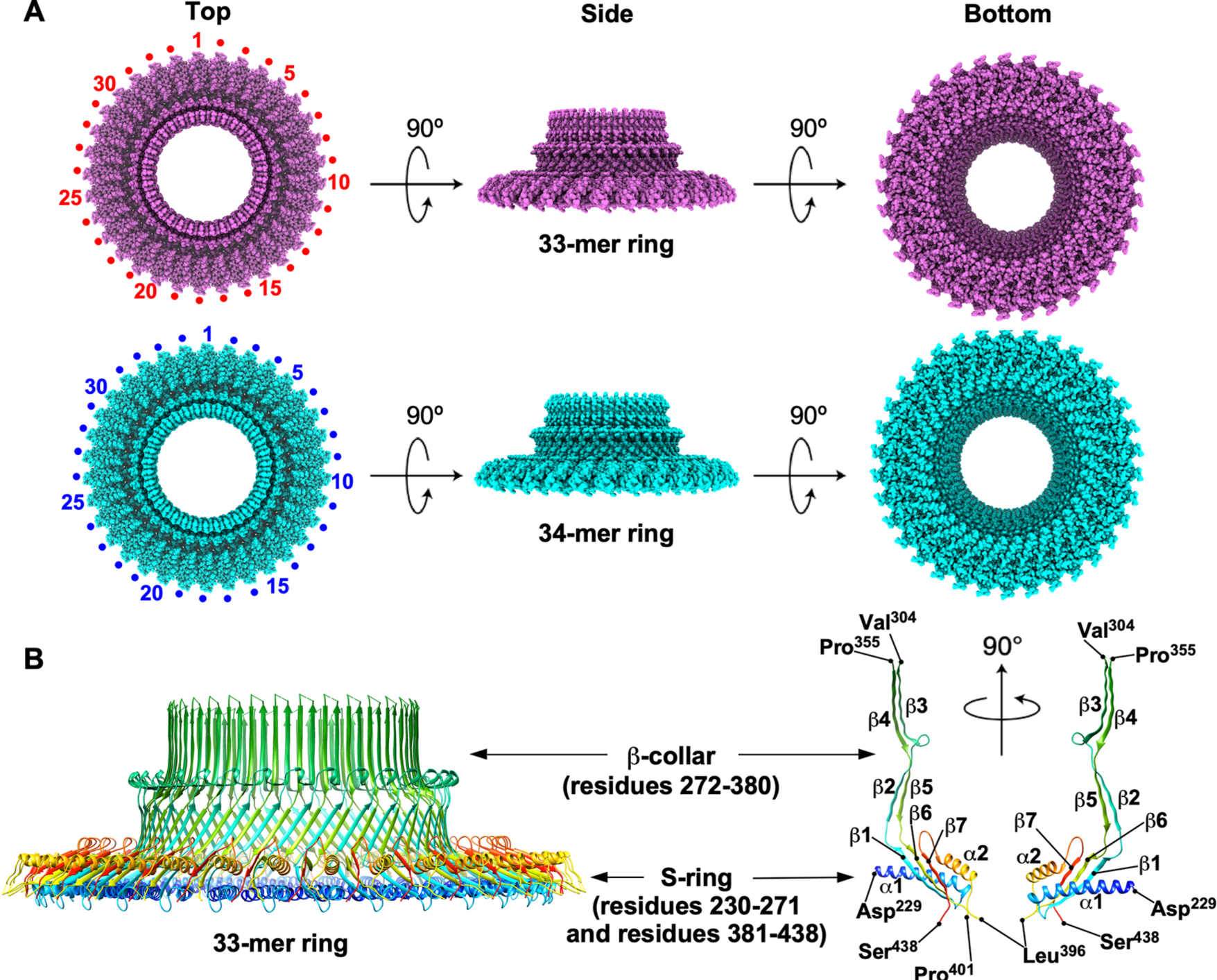
The RBM3-ring structures. **(A)** CryoEM 3D image reconstruction of the 33-mer and 34-mer RBM3-rings with C33 (EMDB ID: EMD-60008) (upper) and C34 (EMDB ID: EMD-60009) (lower) symmetry applied, respectively. **(B)** Cα ribbon diagrams of the atomic model of the RBM3-ring with C33 symmetry applied (left) and the RBM3 subunit forming the S-ring and β-collar (right) (PDB ID: 8ZDT). The Cα backbone is color-coded from blue to red, going through the rainbow colors from the N-terminus to the C-terminus.

The S-ring domain (residues 230–271 and 381–438) is composed of two α-helices (α1 and α2) and three β strands (β1, β6, and β7). Residues 397–400 connecting β6 and α2 are invisible. The three β strands form a core β-sheet, which together with the two α-helices form a globular domain (Fig. 1B). The S-ring domains are horizontally packed with their major axis oriented in the radial direction to form the S-ring. The β-collar domain (residues 272–380) is a long, extended up-and-down β structure consisting of two sets of antiparallel β sheets (β2/β5 and β3/β4) (Fig. 1B). Intermolecular β2-β5 and β3-β4 interactions create a stable and strong β-barrel structure above the S-ring. The chain connecting β3 and β4 at the top of the collar (residues 305–354) is invisible, in agreement with previous reports (5–7,28). The *fliF(N318T)* mutant easily releases the rod-hook-filament structure from the MS-ring under viscous conditions (21), and extragenic suppressor mutations in FlgC or FlgF partially suppress such detachment from the MS-ring (38), suggesting that the N318T mutation affects the interaction between the MS-ring and the proximal rod. Because the N318T substitution is located within the invisible region of the β-collar domain, we suggest that this flexible region may associate with the rod to prevent it from dislodging from the MS-ring when the flagellar motor operates under viscous conditions.

### Structural comparison of the 33-mer and 34-mer RBM3-rings

To clarify why the MS-ring shows a small variation in the ring stoichiometry, we first measured the inner diameters of the 33-mer and 34-mer RBM3-rings (Fig. S5 and Table S2). The inner diameter of the S-ring was 142 Å and 146 Å (as derived from least square fitting of a circle to Cα atoms of Asp-229) and the inner diameter of the β-collar was 101 Å or 105 Å (as derived from least square fitting of a circle to Cα atoms of Pro-355) for the 33-mer and 34-mer rings, respectively. The structures of the S-ring and β-collar domains in the 33-mer and 34-mer rings were nearly identical to each other (Fig. S6A). The subunit interfaces between the S-ring domains in the 33-mer and 34-mer rings were also almost identical (Fig. S6B). When a subunit in the 34-mer ring (labeled A in Fig. S6C) was superimposed on a subunit in the 33-mer ring, misalignment gradually increases as the subunit position moves away from the superimposed ones, reflecting a slightly different curvature of the two rings produced by a subtle difference in the relative angle between the adjacent S-ring domains (Fig. S6C). Interestingly, when the RBM3 structures of previously reported 33-mer and 34-mer rings were superimposed on to those of our 33-mer and 34-mer rings, respectively, with the β-collar domain, subtle differences were observed in the orientation of the S-ring domain relative to the β-collar domain (Fig. S4), producing slight differences in the diameter of the RBM3-ring despite the same rotational symmetry (Table S2). Therefore, we suggest that conformational flexibility of loops connecting with the secondary structures within the RBM3 region produces not only slight differences in curvature of the MS-ring but also produces the MS-ring with different rotational symmetry. This indicates a flexible nature of the loops connecting the β-collar and S-ring domains, which may also be responsible for the small variation in the rotational symmetry of the MS-ring.

### CryoEM structural analysis of the inner part of the M-ring

To build the atomic model of the inner part of the M-ring formed by the RBM1 and RMB2 domains, we performed iterative 3D refinement of the 33-mer ring with C11 symmetry to obtain a higher-resolution map of the inner core ring formed by 22 RBM2 domains. The resolution of the 3D map was improved from 4.2 Å to 3.1 Å (EMDB ID: EMD-60007) (Fig. 2A and Fig. S3C, D), and we were able to build the atomic model of the inner M-ring with a gear wheel-like feature, consisting of the inner core ring formed by 22 RBM2 domains (RBM2_inner_) and 11 cog-like domains, each formed by RBM1 and RBM2, located just outside the inner core ring (PDB ID: 8ZDS) (Fig. 2B). Some blurred densities possibly corresponding to the remaining 22 RBM1 domains were observed around the inner core ring part but was not clear enough to place the atomic model, as reported in the previous studies (11,28). The RBM1 and RBM2 domain structures of *Salmonella* FliF were nearly identical to those of the crystal structure of *Aquifex* FliF (PDB ID: 7CIK) (Fig. S7). However, the relative orientation of RBM1 to RBM2 was distinctly different from that seen in the *Aquifex* FliF crystal structure, suggesting that the domain packing in the crystal was a crystal packing artifact.

**FIG 2.**
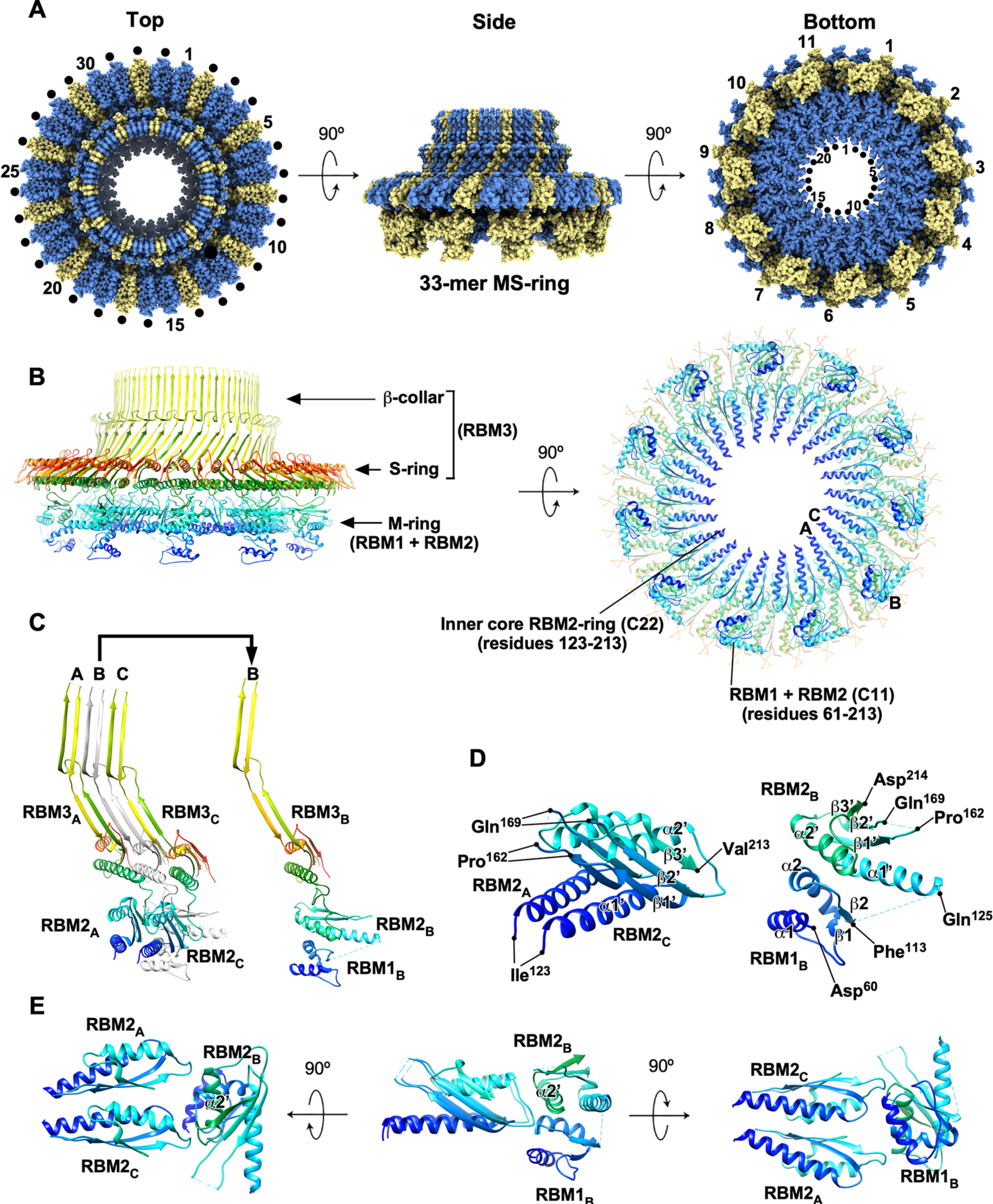
The RBM1-RBM2-RBM3-ring structure revealed by C11 symmetry enforcement. **(A)** CryoEM 3D image reconstruction of the 33-mer MS-ring with C11 symmetry applied (EMDB ID: EMD-60007). All 33 RBM3 domains form the 33-mer S-ring and β-collar, while 22 RBM2 domains face inward to form the inner core ring of the M-ring, and the remaining 11 RBM2 domains form cog-like structures with 11 RBM1 domains on the outer surface of the inner core ring. The densities for the remaining 22 RMB1 domains, presumably located below the 22-mer inner core RMB2-ring, are not observed clearly. FliF subunits are colored either cornflower blue or khaki to identify the two different conformations: cornflower blue for its RBM2 forming the 22-mer inner core ring; and khaki for its RBM2 forming the cog-like structure. **(B)** Cα ribbon diagrams of the atomic model (PDB ID: 8ZDS). The Cα backbone is color-coded from blue to red, going through the rainbow colors from the N-terminus to the C-terminus. RBM2 (residues 123–213) forms the 22-mer inner core ring as well as the 11 cog-like structures with RBM1 (residues 61–113) just outside the inner core ring. **(C)** Three FliF subunits (Mol-A, Mol-B, and Mol-C) of an asymmetric unit of the 33-mer ring, viewed from inside the MS-ring. The RBM2 domains of Mol-A and Mol-C face inward to form the inner core ring. The RBM1 and RBM2 domains of Mol-B form a cog-like structure, in which RBM2 is rotated about 100° in the CCW direction relative to those of Mol-A and Mol-B. **(D)** Interactions between adjacent RBM2 domains in the inner core ring (left panel) and those between RBM1 and RBM2 in the cog-like structure (right panel). **(E)** Three different views of RBM1-RBM2 of Mol-B in the cog-like structure interacting with 2 RBM2 domains of Mol-A and Mol-C forming the 22-mer inner core ring on the left.

FliF showed two distinct conformations in the 33-mer MS-ring as reported previously (28,29). The asymmetric unit of the 33-mer ring, for which the 3D map was reconstructed with C11 symmetry, consists of 3 FliF molecules, Mol-A, Mol-B, and Mol-C. Mol-A and Mol-C have RBM2 (residues 123–213) and RBM3 while Mol-B has RBM1 (residues 61–113) in addition to these two domains (Fig. 2C). RBM1 is composed of two α-helices (α1 and α2) and a parallel two-stranded β-sheet (β1 and β2). RBM2 consists of two α-helices (α1’ and α2’) and an antiparallel three-stranded β-sheet (β1’– β3’). Residues 163–168 of RBM2 are not visible (Fig. 2D). Although residues 114–124 connecting RBM1 and RBM2 are invisible in Mol-B, RBM1 tightly associates with RBM2, thereby stabilizing the RBM1 domain structure in Mol-B (Fig. 2D, right panel). Extensive RBM2-RBM2 intersubunit interactions produce a flat inner core ring of the M-ring (Fig. 2B, left panel). The RBM1-RBM2 domains of Mol-B is oriented as its RBM2 rotated about 100° in the CCW direction relative to RBM2 of Mol-A and Mol-C forming the inner core ring and is bound to the outer surface of the inner core ring (Fig. 2E). Interestingly, RBM1 binds to two RBM2 domains (Fig. 2E, left and right panels), creating a discrete cog-like structure just outside the inner core ring (Fig. 2A, middle and right panels, and Fig. 2B).

### Mutational analysis of the i-loop of RBM2

The i-loop connecting β1’ and β3’ of the RBM2 domain, which consists of residues 159–172, extends toward the central hole of the inner core ring. In-frame deletion of residue 161–170 in FliF cause a loss-of-function phenotype, suggesting that the i-loop is critical for FliF function (29). We found that residues 163–168 in the i-loop are disordered and invisible in the MS-ring structure obtained in this study (Fig. S8, right panel) as well as in the 6SCN and 8T8P structures (11,28). In contrast, the i-loop is well ordered in the native basal body MS-ring (PDB ID: 7NVG) possibly by direct contact with FliP and FliR of the export gate complex located in the central pore (5,6) (Fig. S8, left panel). Therefore, we investigated whether the contact of i-loop with FliP or FliR is critical for stable incorporation of the export gate complex into the central pore. To do so, we constructed two *fliF* deletion mutants lacking either residues 164–167 or residues 165–167 and analyzed their motility in soft agar (Fig. S9A). Immunoblotting with polyclonal anti-FliF antibody revealed that these two deletions did not affect the steady cellular level of FliF at all (Fig. S9B). Wild-type FliF fully restored motility of a Δ*fliF* mutant. The Δ164–167 and Δ165–167 mutant variants did not complement the Δ*fliF* mutant at all (Fig. S9A).

The MS-ring is the housing for the transmembrane export gate complex of the fT3SS. Therefore, we tested whether the lack of motility of the Δ164–167 and Δ165–167 *fliF* mutants is a consequence of the lack of protein export activity of the fT3SS. We analyzed the secretion level of FlgD as a representative export substrate of the fT3SS by immunoblotting with polyclonal anti-FlgD antibody and found that these two deletions inhibit FlgD secretion (Fig. S9B). The deletion of residues 165–167 did not inhibit MS-ring formation (Fig. S9C), suggesting that these deleted residues are important for stable interactions of the i-loop with FliP or FliR, thereby allowing the export gate complex to be efficiently accommodated inside the central pore of the inner core ring.

### Mutational analysis of the RBM2-3 loop connecting RBM2 and RBM3

The RBM2-3 loop connecting RBM2 and RBM3 (residues 214–228) is highly flexible, allowing FliF to adopt two different conformations in the MS-ring (Fig. 3A, upper panels). Furthermore, conformational flexibility of this loop possibly allows the position, orientation, and curvature of the inner core ring to be adjusted, thereby causing subtle structural differences in this ring with 11 cog-like structures formed by RBM1-RBM2 attached on its outer surface (Fig. S10). The RBM2-3 loop contains a conserved DQxGxxL motif (residues 214–220), in which the third and fifth residues are commonly hydrophilic (Fig. 3A, lower panels). To clarify the role of the DQxGxxL motif in flagellar assembly, we deleted this motif from FliF and analyzed the effect on motility in soft agar (Fig. 3B). We also constructed the *fliFΔ*(*221–227*) mutant (Fig. S11A). Deletion of residues 214–220 or residues 221–227 did not affect the steady cellular level of FliF (Fig. 3C and Fig. S11B). Deletion of residues 214–220 resulted in a non-motile phenotype (Fig. 3B), and FlgD and FlgL were not secreted into the culture media (Fig. 3C). Because this deletion did not inhibit MS-ring formation (Fig. 3D), we suggest that deletion of the DQxGxxL motif inhibits the assembly of export gate complex into the central pore of the inner core ring. On the other hand, the *fliFΔ*(*221–227*) mutant displayed a weak motile phenotype (Fig. S11A). Consistently, secreted FlgD and FlgL were present in the culture media albeit lower than the wild-type levels (Fig. S11B). Since deletion of residues 221–227 did not inhibit MS-ring formation (Fig. S11C), we suggest that an appropriate length of the RBM2-3 loop is required for efficient and robust incorporation of the export gate complex into the MS-ring.

**FIG 3.**
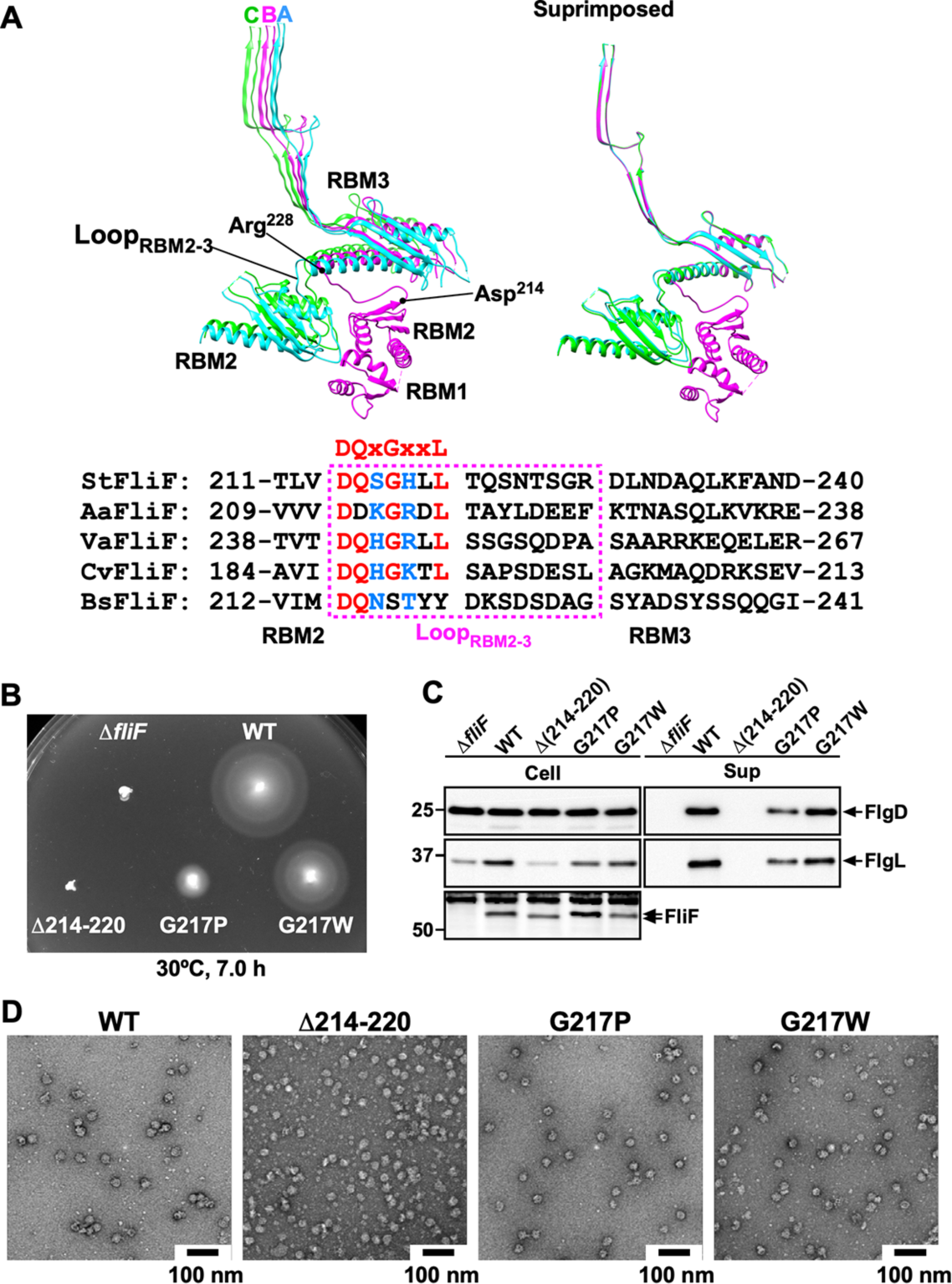
Mutational analysis of the RBM2-3 loop. **(A)** Structural comparison and multiple sequence alignments of the RBM2-3 loop. Three FliF subunits, Mol-A, Mol-B and Mol-C of the asymmetric unit of the 33-mer MS-ring are shown in the left panel. The RBM2 domains of Mol-A and Mol-C (green) form part of the inner core ring, and RBM1-RBM2 of Mol-B (magenta) binds to the outer surface of the inner core ring with its RBM2 rotated about 100° relative to those forming the inner core ring. The three subunits were superimposed with β-collar in the right panel. Multiple sequence alignment was carried out by Clustal Omega. Conserved residues are highlighted in red. The third and fifth residues in the conserved DQxGxxL motif are commonly hydrophilic. UniProt Accession numbers: *Salmonella* (StFliF), P15928; *Aquifex* (AaFliF), A0A7C5L200; marine *Vibrio* (VaFliF), A0A1W6UMQ1; *Caulobacter* (CvFliF), Q04954; *Bacillus* (BsFliF), P23447. **(B)** Motility of a *Salmonella fliF* null mutant harboring pTrc99AFF4 (indicated as Δ*fliF*), pMKMiF015 (indicated as WT), pMKMiF015(Δ214–220) (indicated as Δ214–220), pMKMiF015(G217P) (indicated as G217P), or pMKMiF015(G217W) (indicated as G217W) in soft agar. The plate was incubated at 30°C for 7 hours. **(C)** Flagellar protein secretion assays. Whole cell proteins (Cell) and culture supernatant fractions (Sup) were prepared from the above transformants. A 5 μl solution of each protein sample, which was normalized to an optical density of OD_600_, was subjected to SDS-PAGE, followed by immunoblotting with polyclonal anti-FlgD (first row), anti-FlgL (second row) or anti-FliF (third row) antibody. The positions of molecular mass markers (kDa) are shown on the left. **(D)** Negative stained EM images of the MS-rings isolated from the above transformants.

To further clarify the role of the DQxGxxL motif, we constructed 8 point mutants, *fliF(D214A)*, *fliF(D214E)*, *fliF(D214R)*, *fliF(G217A)*, *fliF(G217P)*, *fliF(G217W)*, *fliF(L219A)*, and *fliF(L220A)*, and analyzed their motility in soft agar (Fig. 3B and Fig. S11A). No FliF was seen in the *fliF(D214R)* mutant while FliF was detected at the wild-type level in the remaining mutants (Fig. 3C and Fig. S11B). When Arg residues are exposed on the surface of proteins in structurally flexible regions, their C-terminal side is easily cleaved by Arg-C endoprotease. The destabilization of FliF by the D214R mutation (Fig. S11B) suggests a highly flexible conformation of the DQxGxxL chain. The D214A, D214E, L219A, and L220A mutant variants complemented the Δ*fliF* mutant to the wild-type level, indicating that the Arg-214, Leu-219, and Leu-220 residues are not so important (Fig. S11A). The G217A and G217W mutations slightly reduced motility, and the G217P mutation inhibited motility (Fig. 3B and Fig. S11A). Consistently, the G217A and G217W mutations reduced the levels of FlgD and FlgL secretion slightly, and the G217P mutation did so significantly (Fig. 3C and Fig. S11B). Because neither G217A, G217P, nor G217W substitution affected MS-ring formation (Fig. 3D and Fig. S11C), the conformational flexibility of the DQxGxxL motif is important for creating a hole of an appropriate size and form in the inner core ring to accommodate the export gate complex.

### Isolation of up-motile mutants from the *fliF(I252A)* mutant

Ile-252 of RBM3 is located at an interface between adjacent FliF subunits in the MS-ring and critical for the FliF function (Fig. S6B) (7). The *fliF(I252A)* mutation does not affect the formation of the MS-C-ring complex *in vivo*, but significantly reduces the flagellar protein export activity (7), raising the question of how this mutation affects the assembly of fT3SS into the base of the flagellum. To clarify this question, we isolated eight up-motile mutants form the *fliF(I252A)* mutant (Fig. 4A). DNA sequencing identified two missense mutations, A252V (isolated four times) and R228C (isolated four times) in FliF (Fig. 4B). The motility of the *fliF(A252V)* mutant was much better than that of the *fliF(I252A)* mutant although not as good as the wild-type strain. Because Ile-252 and Ile-256 make hydrophobic contacts with Val-390 in their neighboring FliF subunit, stabilizing the RBM3-ring, the length of hydrophobic side chain of Ile-252 seems important for the intermolecular RBM3-RBM3 interactions. On the other hand, the R228C mutation is located at the C-terminus of the RBM2-3 loop in Mol-A and Mol-C and at the N-terminus of the α1 helix of RBM3 in Mol-B (Fig. 4B). This suggests that the I252A substitution affects the orientation of the RBM2-3 loop relative to RBM3. Therefore, the R228C mutation would induce a conformational change of this loop, allowing the inner core ring to form an appropriate central hole for the export gate complex to efficiently assemble at the center of the MS-ring even in the presence of the I252A mutation.

**FIG 4.**
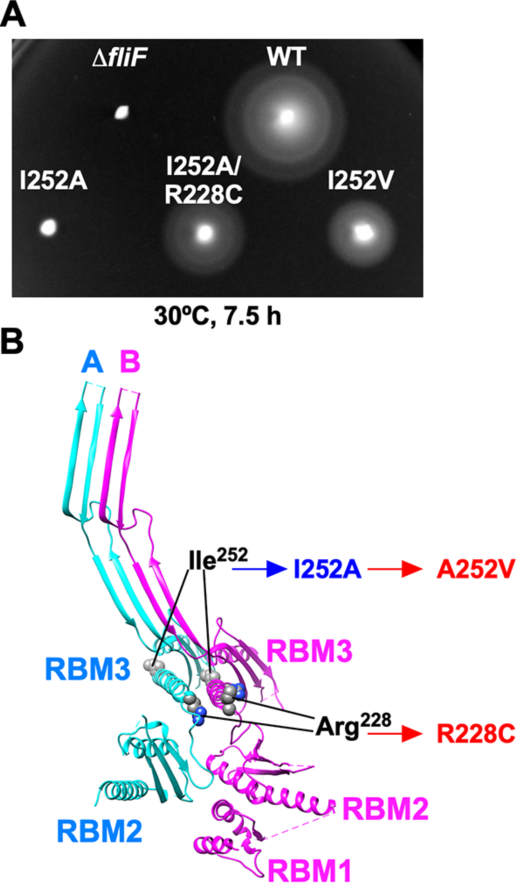
Isolation of gain-of-function mutants from the *fliF(I252A)* mutant. **(A)** Motility of a *Salmonella fliF* null mutant harboring pET3c (indicated as Δ*fliF*), pMKMiF001 (indicated as WT), pMKMiF002 (indicated as I252A), pMKMiF002-SP1 (indicated as I252A/R228C), or pMKMiF002-SP2 (indicated as I252V) in soft agar. The plate was incubated at 30°C for 7.5 hours. **(B)** Location of intragenic suppressor mutations isolated from the *fliF(I252A)* mutant. Two FliF subunits (Mol-A and Mol-B) are viewed from the inside the MS-ring. The I252A mutation and its suppressor mutations are highlighted in blue and red, respectively.

## DISCUSSION

FliF is folded into two different conformations to build the MS-ring with three different, C34, C23, and C11 symmetries in different structural parts to exert its multiple functions (5–7,27,28). However, the self-assembly mechanism of the MS-ring was not fully understood. Therefore, we performed high-resolution cryoEM image analysis of the MS-ring formed by FliF co-expressed with FliG and transmembrane export gate proteins and provided evidence that the intermolecular RBM3-RBM3 interactions generate two distinct orientations of RBM2 relative to RBM3 through a conformational change of the RBM2-3 loop connecting RBM2 and RBM3 to form the inner core ring with its central hole of an appropriate size for the export gate complex to efficiently assemble there as well as the 11 cog-like structures formed by RMB1-2 just outside the inner core ring.

Although RMB3 has a stably folded domain structure, a subtle change in its conformation still allows FliF to form the MS-ring with different rotational symmetries due to a subtle variation in the ring curvature during FliF assembly (Figs S5 and S6 and Table S2). The MS-ring formed by co-expression of full-length FliF with FliG, FliM, and FliN has the rotational symmetry ranging from 32-fold to 34-fold, with 33-mer rings comprising most particles (80% of the population) (36). In contrast, we found that the MS-ring structure showed two distinct classes, 33-mer (53% of the population) and 34-mer (47% of the population), when FliF was co-expressed with FliG and export gate proteins (Fig. S3B). Because the MS-ring in the native basal body has only C34 symmetry, we propose that the export gate complex also contributes to the precise determination of the MS-ring curvature, thereby creating only the 34-mer MS-ring in the basal body.

The flagellar export gate complex is accommodated in the central hole of the MS-ring. Each i-loop of 22 RBM2 domains forming the inner core ring directly makes a contact with FliP or FliR in the native basal body MS-ring (Fig. S8, left panel). Here, we showed that deletion of residues 165–167 involved in this contact inhibited flagellar protein export but not MS-ring formation (Fig. S9). We also found that residues 163– 168 in the i-loop are invisible when the FliP_5_-FliQ_4_-FliR_1_ complex is missing (Fig. 2D and Fig. S8, right panel). Therefore, we suggest that the interaction of the i-loop with FliP or FliR stabilizes the i-loop conformation, allowing the FliP_5_-FliQ_4_-FliR_1_ complex to firmly bind to the MS-ring. The RBM1 domains below the inner core ring are invisible even in the isolated native basal body containing FliP, FliQ, and FliR. Because FlhA and FlhB are missing in the isolated native basal body, RBM1 may bind to these two export gate proteins to be stabilized.

The RBM2-3 loop of FliF contains a relatively well-conserved DQxGxxL motif (Fig.3A). Deletion of this motif resulted in a non-motile phenotype, indicating the critical importance of this motif (Fig.3). Furthermore, mutational analysis of this motif revealed that a conformational flexibility of the conserved Gly-217 residue is important (Fig.3). Therefore, we propose that the conformational flexibility of the DQxGxxL motif allows the inner core ring to form an appropriate central hole for the export gate complex to efficiently assemble at the center of the MS-ring.

How does FliF form the MS-ring with three different rotational symmetries within its ring structure? The *Salmonella* FliF fragment containing RBM1, RBM2, and RBM3 forms the inner core ring, S-ring and β-collar, suggesting that two transmembrane helices of FliF are not involved in MS-ring formation. The RBM3 fragment alone can form the S-ring and β-collar, suggesting that neither RBM1 nor RBM2 is required for RBM3-ring formation. The RBM2 fragment alone can self-assemble into the inner core ring while the RBM1-RBM2 fragment cannot. Because the purified RBM1-RBM2 fragment exits as a stable monomer in solution, this suggests that RBM1 prevents RBM2_-_ring formation (36). Therefore, in full-length FliF assembly, intermolecular RBM3-RBM3 interactions first promote the initiation of RBM3-ring formation, and then it is followed by the formation of the inner core RBM2-ring and the cog-like structures just outside the ring.

In our present MS-ring structure, we found that RBM1 binds to the α1’ and α2’ helices of RBM2 in the cog-like structure (Fig. 2D, right panel). Leu-98 of RBM1 makes hydrophobic contacts with Leu-147, Ala-193, and Leu-197 of RBM2 (Fig. 5A, left panel). However, these two α-helices of RBM2 and its residues involved in the interaction with RMB1 are also involved in RBM2-ring formation. Extensive intermolecular interactions between these two α-helices of RBM2 and an antiparallel β-sheet of its neighboring RBM2 as well as intermolecular α1’-α1’ interactions promote RBM2-ring formation (Fig. 2D, left panel). In the RBM2-ring, Val-213 of RBM2 of Mol-A makes hydrophobic contacts with Leu-147, Ala-193, and Leu-197 of RBM2 of Mol-C (Fig. 5A, right panel). Furthermore, Phe-126, Gln-133, and Arg-134 of RBM2 of Mol-A interact with Gln-129, Tyr-132, Glu-139 of RBM2 of Mol-C, respectively. Because the density corresponding to RBM1 was not clearly observed below the RMB2-ring, it seems likely that RBM1 dissociates from RBM2 during RBM2-ring formation. Because the RBM1-RBM2 fragment forms a helical tubular structure along with the RBM3 fragment *in vitro* (36), RBM1 would bind to the oligomerization surface of RBM2 before the next FliF subunit binds, thereby preventing uncontrolled MS-ring formation. Based on this idea, we built a structural model of the monomeric RBM1-RBM2-RBM3 structure (RBM1-3) by fitting the RBM1-RBM2 structure (RMB1-2) on an RBM2-RBM3 unit in the 33-mer MS-ring using RBM2, followed by connecting RBM3 with the fitted RBM1-2 at Leu-219 (Fig. 5B). When the two RBM1-3 models are placed side by side by fitting their RBM3 domains to those in the 33-mer MS-ring, serious steric hinderance occurs between RBM1 and RBM2 (Fig. 5C, open arrow) or between the RBM1-2 domains of each subunit (Fig. 5D). Because RBM1-2 of one FliF subunit is bound to the outside of ring-forming RBM2 domains of two closest FliF subunits (Fig. 2E), the RBM1-2 units of the two FliF subunits move around each other at the initial stage of FliF oligomerization, and finally the RBM1-2 unit of one subunit weakly associates with RBM2 of the other subunit as seen in Mol-B and Mol-C in the 33-mer MS-ring (Fig. 5E).

**FIG 5.**
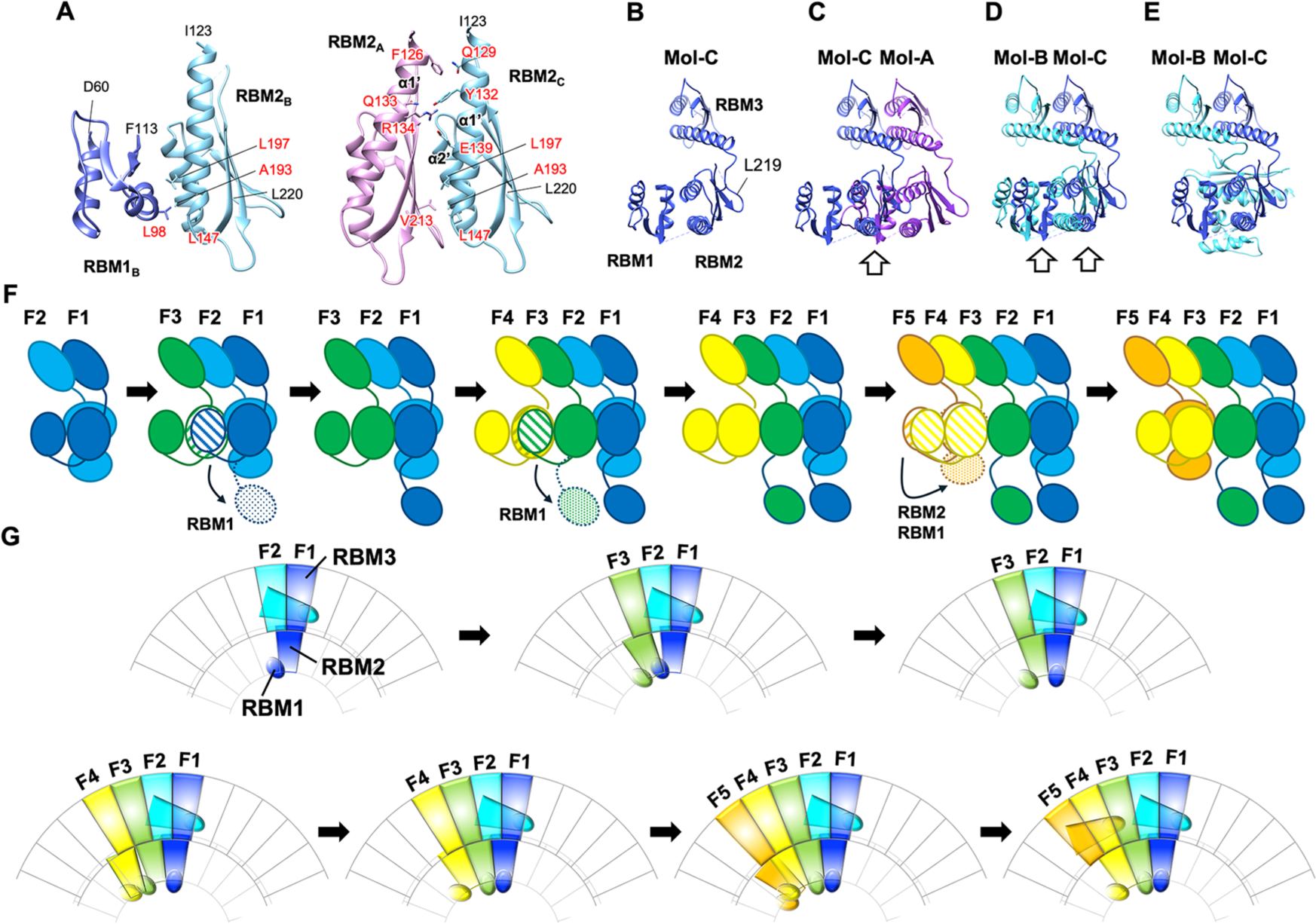
Model for MS-ring formation with three different symmetries. **(A)** Side-by-side comparison of RBM1-RBM2 in the cog-like structure (left panel) and RMB2-RMB2 forming the inner core ring (right panel) shows that both use the same surface of RBM2 for complex formation. The residues involved in the interactions are indicated by labels in red. **(B)** A plausible structural model of RBM1-RBM2-RBM3 (RBM1-3) as an assembly unit for MS-ring formation. **(C)** When two RBM1-3 units of Mol-A (violet) and Mol-C (blue) start forming the MS-ring, RBM1 of Mol-A collides with RMB2 of Mol-C (indicated by open arrow). **(D)** When two RBM1-3 units of Mol-B (cyan) and Mol-C (blue) form a dimer for initiating MS-ring formation, RBM1-2 of Mol-C on the right collides with RMB1-2 of Mol-B on the left (indicated by open arrow). **(E)** A possible structural model for the initial FliF dimer for MS-ring formation is two RBM1-3 units of Mol-B (cyan) and Mol-C (blue) with RBM1-2 of Mol-B binding to the outer surface of RBM2 of Mol-C forming the inner core ring. **(F, G)** The CCW growth model of MS-ring formation viewed from the center **(F)** and periplasmic side **(G)** of the MS-ring. After forming the initial FliF dimer (F1 and F2) (step 1), the third subunit (F3) binds to it, but steric hinderance occurs between RBM2 of F3 and RBM1 of F1 (shown by stripe pattern), thereby inducing the dissociation of RBM1 from RBM2 in F1 and its move to a space below RBM2 (shown by hatched pattern) (step 2). This allows RBM2 of F3 to firmly associate with RBM2 of F1 for initiating inner core ring formation, with RBM1-2 of F2 strongly associated with these two RBM2 units on their outer surface to form a cog-like structure (step 3). The fourth subunit (F4) binds to F3, but steric hinderance between RBM2 of F4 and RBM1 of F3 (shown by stripe pattern) induces the dissociation of RBM1 from RBM2 in F3 and its move to a space below RBM2 of F3 (shown by hatched pattern) (step 4), allowing RBM2 of F4 to firmly associate with RBM2 of F3 (step 5). When the fifth subunit (F5) binds to F4, steric hinderance occurs between two RBM1-2 units of F5 and F4 (shown by stripe pattern) (step 6). To avoid this collision, RBM1-2 of F5 moves outward through a large conformational change of the RMB2-3 loop connecting RBM2 and RBM3 and binds to the outer surface of the RBM2 domains of F3 and F4 to from the second cog-like structure while RBM1 of F4 remains associated with its RBM2 (step 7). The FliF assembly process will proceed by repeating these steps.

Assuming a FliF dimer as the initial complex for MS-ring formation, we propose a model of the MS-ring assembly process as follows (Fig. 5F, G). The schematic figures explaining the assembly process presented in Fig. 5F and 5G are viewed from the center and periplasmic side of the MS-ring. After the initial dimer is formed (Step 1), RMB3 of the third FliF subunit (F3) binds to RMB3 of F2 on the left side. However, due to a steric hindrance, RBM1-2 of F3 cannot be placed to the left (Fig. S12B) or outside (Fig. S12C) of RBM2 of F1. Therefore, RBM1-2 of F3 moves closer to RBM1 of F1 and competes with it to bind to the left surface of RBM2 of F1. Since the intermolecular RBM2-RMB2 interaction is more extensive than the interdomain interaction between RBM1 and RBM2 (Fig. 5A), RBM1-2 of F3 induces the dissociation of RBM1 from RBM2 in F1 (Fig. 5F, G, step 2), and RBM2 of F3 can now associate with RBM2 of F1 to start forming the RBM2-ring (Step 3). Extensive RBM2-RBM2 interactions results not only in the initiation of RBM2-ring formation but also in the stabilization of the first RBM1-2 cog-like structure just outside the two ring-forming RBM2 domains (Fig. S12D). Then, RBM3 of the fourth FliF subunit (F4) binds to the left surface of RBM3 of F3 (Fig. 5F, G, step 4). However, again due to a steric hindrance, RBM1-2 of F4 cannot be placed to the left of RBM2 of F3 (Fig. S12E) or outside the RBM2-ring (Fig. S12F). Thus, RBM1-2 of F4 replaces RBM1 of F3 (Fig. 5F, G, step 5 and Fig. S12G), just as when F3 bound to the initial dimer. Then, RMB3 of the fifth FliF subunit (F5) binds to the left surface of RMB3 of F4 (Fig. 5F, G, step 6). Because of a steric hinderance, RBM1-2 of F5 cannot bind to the left of RBM2 of F4. However, because the RBM2 domains of F3 and F4 already formed a stable docking site for RBM1-2 of F5, it can rotate about 100° relative to the RBM2 domains of F3 and F4 to form the second cog-like unit outside the RBM2-ring (Step 7). By repeating these processes, the FliF assembly grows in the CCW direction when viewed from the periplasm, and each RBM1-2 is placed outside the RBM2-ring to form the cog-like structure for every three FliF subunit assembly.

Finally, when the 33rd subunit (F33) is inserted between F32 and F1, the RBM1 domains of F32 and F33 dissociate from RBM2, thereby not only closing the RBM2-ring with C22 symmetry but also stabilizing the last of the 11 cog-like structures. The 33-mer MS-ring with three different symmetries, C33, C22, and C11, is thus completed (Fig. S13A). The 34-mer MS-ring can be formed through the same process except for the final step, which is slightly different from that in 33-mer ring formation (Fig. S13B).

In the above model, MS-ring formation proceeds in the CCW direction, but it might also proceed in the CW direction (Fig. S14). In the CW growth model, RMB3 of F3 binds to RMB3 of F1 in the initial dimer on the right side (Fig. S14A, B). Due to a steric hindrance, RBM1-2 of F3 cannot be placed on the right side of RBM2 of F1 but can be placed outside the ring-forming RBM2 domains (Fig.S14B). Unlike the initial dimer formation, RBM1-2 of F3 cannot be stabilized on the outside the ring by binding to RBM2 of other subunits (Fig.S14C). RBM1-2 of F3 moves freely, and no further growth of FliF assembly would occur. Therefore, we suggest that the CCW growth model is much more plausible than the CW growth model.

## MATERIALS AND METHODS

### Bacterial strains, plasmids, and DNA manipulations

Bacterial strains and plasmids used in this study are listed in Table S3. The pMKM2001 plasmid encodes FliO, His-FliP, HA-FliQ,FliR-FLAG, FlhA, FlhB, FliF, and FliG. FliO, His-FliP, HA-FliQ,FliR-FLAG, FlhA, and FlhB are expressed from a p*trc* promotor whereas FliF and FliG are expressed from an arabinose-inducible promotor. DNA manipulations were performed using standard protocols. Site-directed mutagenesis was carried out using Prime STAR Max Premix as described in the manufacturer’s instructions (Takara Bio). All *fliF* mutations were confirmed by DNA sequencing (Eurofins Genomics).

### Expression and purification of the MS-ring

A 13 ml of overnight culture of *Salmonella* SJW1368 [Δ(*cheW-flhD*)] cells harboring pMKM20001 (pTrc99CES3/ FliO + His-FliP + HA-FliQ + FliR-FLAG + FlhA + FlhB + FliF + FliG) were inoculated into a 1.3 l of fresh 2×YT [1.6% (w/v) Bacto-tryptone, 1.0% (w/v) Bacto-yeast extract, 0.5% (w/v) NaCl] containing 100 µg mL^−1^ ampicillin, and the cells were grown at 30°C until the cell density had reached an OD_600_ of about 0.6. After 30 min incubation at 4°C, arabinose was added at a final concentration of 0.2%, and the cells were grown at 16°C for 12 hours. The SJW1368 cells co-expressing FliF, FliG, and the transmembrane export gate complex were collected by centrifugation (6,400 × *g*, 10 min, 4°C) and stored at −80°C. The cells were thawed, resuspended in 80 ml of 50 mM Tris-HCl, pH 8.0, 5 mM EDTA, 50 mM NaCl and disrupted using a French press at a pressure level of 8,000 psi (FA-032, Central Scientific Commerce). After cell debris and undisrupted cells were removed by centrifugation (20,000 × *g*, 20 min, 4°C), crude membranes were isolated by ultracentrifugation (90,000 × *g*, 1 h, 4°C). The harvested membranes were solubilized in 44 ml of 50 mM CAPS-NaOH, pH 11.0, 50 mM NaCl, 5 mM EDTA, 1% (w/v) Triton X-100 and left on ice for 1 hour. After centrifugation (20,000 g, 20 min, 4°C), the supernatant was ultracentrifuged (90,000 × *g*, 1 h, 4°C), and the pellet was resuspended in 3.8 ml of 25 mM Tris-HCl, pH 8.0, 50 mM NaCl, 1 mM EDTA, 0.1% (w/v) Triton X-100, 0.05% (w/v) lauryl maltose neopentyl glycol (LMNG), and the sample was loaded onto a 15–40% (w/w) sucrose density gradient. After ultracentrifugation (49,100 × *g*, 13 h, 4°C), fractions containing FlhA, FliF, and FliG were collected (Fig. S2) and concentrated by ultracentrifugation (90,000 × *g*, 1 h, 4°C). The pellet was resuspended in 20 μl of 50 mM Tris-HCl, pH 8.0, 50 mM NaCl, 25 mM imidazole, 0.05% (w/v) Triton X-100, 0.05% (w/v) LMNG.

### Sample vitrification and cryoEM data acquisition

Quantifoil Cu 200 mesh R0.6/1.0 holey carbon grids (Quantifoil) were glow discharged on a glass slide for 20 sec. A 2.7 µl aliquot of the sample solution was applied to the grid and blotted by a filter paper for 3 sec at 100% humidity and 4°C. The grids were quickly frozen in liquid ethane using Vitrobot IV system (Thermo Fisher Scientific). The grids were inserted into a CRYO ARM 300 transmission electron microscopy (JEOL Ltd. Japan) equipped with a cold field-emission electron gun operated at 300 kV and an Ω-type energy filter with a 20^−^ eV slit width. CryoEM images were recorded with a K3 direct electron detector camera (Gatan, USA) at a nominal magnification of ×50,000, corresponding to an image pixel size of 1.0 Å, using SerialEM (39). The holes were detected using YoneoLocr (40). Movie frames were recorded in CDS counting mode with a total exposure time of 3 sec and a total dose of ∼40 electrons Å^−2^. Each movie was fractionated into 40 frames. In total, 4,885 movies were collected.

### Image processing of the MS ring

Single particle analyses of the MS-ring were performed using Relion-3.1 (41). Image processing procedures of the MS-ring are described in Fig. S3. After performing motion corrections to align all micrographs, followed by the estimation of parameters of the contrast transfer fraction (CTF), particle images were automatically selected via LoG auto-picking, and the selected particles were extracted into a box of 512 × 512 pixels (1,015,741 particles). Particle images from good 2D class average images were selected for the initial 3D model. In total, 757,555 particles were subjected to 3D classification with C1 symmetry into five classes. A 3D class with good MS ring geometry was further divided into three classes with C1 symmetry; two more iteration of the 3D classification resulted in two different good 3D classes. One had C33 symmetry, and the other had C34 symmetry. After 3D refinement for each of these two classes, postprocessing yielded the 3D maps of the 33-mer and 34-mer rings with resolutions of 4.2 Å (216,718 particles) and 4.6 Å (194,233 particles), respectively, according to 0.143 criterion of the Fourier shell correlation (FSC). Then the C33 or C34 symmetry was applied for further image analysis to obtain much the 3D maps with much higher resolutions. After three rounds of CTF refinement and polish, the 3D maps of the 33-mer and 34-mer RBM3-rings were obtained at 2.4 Å and 2.5 Å resolutions, respectively. To obtained detailed structural information on the inner and middle parts of the M-ring, C11 symmetry was applied to the 33-mer ring structure after several iterations of the 3D classification from particles with C33 symmetry. The map of the MS ring with C11 symmetry was obtained at 3.1 Å resolution (68,598 particles). The cryoEM density maps were deposited into Electron Microscopy Data Bank with accession codes EMD-60007 for MS ring with C11 symmetry applied, EMD-60008 for RBM3-ring with C33 symmetry applied, and EMD-60009 for RBM3-ring with C34 symmetry applied.

### Model building and refinement of the MS ring structure

The atomic models of the RBM3-ring with C33 or C34 symmetry and the MS-ring with C11 symmetry were constructed using Coot (42). PHENIX was used for real-space refinement based on the cryoEM maps (43). The refinement statics are summarized in Table S1. The atomic coordinates of the FliF complex have been deposited in the Protein Data Bank with accession codes 8ZDS for MS-ring with C11 symmetry applied, 8ZDT for RBM3-ring with C33 symmetry applied, 8ZDU for RBM3-ring with C34 symmetry applied.

### Motility assays in soft agar

Fresh transformants were inoculated into soft agar plates [1.0% (w/v) Bacto-tryptone, 0.5% (w/v) NaCl, 0.35% (w/v) agar] containing 100 μg/ml ampicillin and incubated at 30°C. The assay was performed at least five times to confirm the reproducibility of the results.

### Secretion assay

A 100 μl of the overnight culture of *Salmonella* cells was inoculated into a 5 ml of fresh L-broth [1% (w/v) Bacto-tryptone, 0.5% (w/v) Bacto-yeast extract, 0.5% (w/v) NaCl] containing 100 μg/ml ampicillin and incubated at 30 °C with shaking until the cell density had reached an OD_600_ of ca. 1.4–1.6. Details of sample preparations have been described previously (44). Both whole cellular proteins and culture supernatants were normalized to a cell density of each culture to give a constant cell number. After sodium dodecyl sulfate-polyacrylamide gel electrophoresis (SDS-PAGE), immunoblotting with polyclonal anti-FlgD, anti-FlgL, or anti-FliF antibody as the primary antibody and anti-rabbit IgG, HRP-linked whole Ab Donkey (GE Healthcare) as the secondary antibody was carried out using iBind Flex Western Device as described in the manufacturer’s instructions (Thermo Fisher Scientific). Detection was performed with Amersham ECL Prime western blotting detection reagent (Cytiva). Chemiluminescence signals were captured by a Luminoimage analyzer LAS-3000 (GE Healthcare). All image data were processed with Photoshop (Adobe). At least three independent experiments were performed.

### Data availability

The cryoEM maps have been deposited in the Electron Microscopy Data Bank under accession code EMD-60007, EMD-60008 and EMD-60009 for the MS-ring with C11, C33, and C34 symmetry applied, respectively. The atomic coordinates for the MS-ring with C11, C33, and C34 symmetry applied have deposited in the Protein Data Bank under accession code 8ZDS, 8ZDT, 8ZDU, respectively.

## Acknowledgements

We thank Kelly T. Hughes for his kind gift of a *Salmonella fliF* null mutant and Yasuyo Abe and Yoshie Kushima for technical assistance. This work was supported in part by JSPS KAKENHI Grant Numbers JP20K15749 and JP22K06162 (to M.K.) and JP19H03182, JP22H02573, and JP22K19274 (to T.Minamino) and MEXT KAKENHI Grant Numbers JP20H05532, and JP22H04844 (to T.Minamino). This work has also been supported by Platform Project for Supporting Drug Discovery and Life Science Research (BINDS) from AMED under Grant Number JP19am0101117 and JP21am0101117(to K.N.), by the Cyclic Innovation for Clinical Empowerment (CiCLE) from AMED under Grant Number JP17pc0101020 (to K.N.), and by JEOL YOKOGUSHI Research Alliance Laboratories of Osaka University (to K.N.).

## Author Contributions

T.Minamino and K.N. conceived and designed research; M.K. prepared samples for cyroEM; M.K., F.M., and T.Miyata collected and analysed cryoEM image data; M.K. and K.I. built atomic models; M.K. and T.Minamino performed genetic, biochemical, and physiological experiments; T.Minamino, K.I., and K.N. wrote the paper based on discussion with other authors.

## Competing interests

The authors declare no competing interests.

## Notes

### Competing Interest Statement

The authors have declared no competing interest.

