## Supplemental information file for "Structural insights into the assembly mechanism of the flagellar MS-ring with three different symmetries"

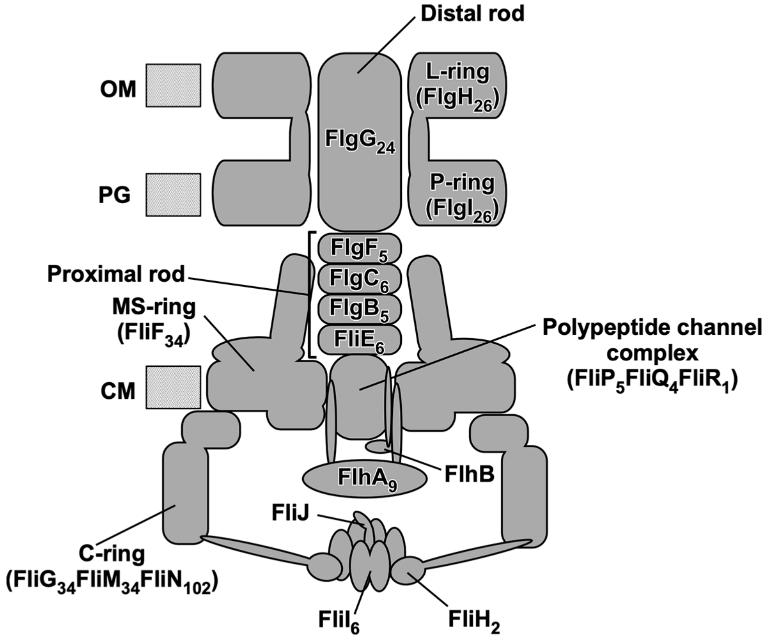

**FIG S1 Schematic diagram of the flagellar basal body.** The flagellar basal body consists of the C-ring (FliG, FliM, FliN), MS-ring (FliF), P-ring (FlgI), L-ring (FlgH) and rod (FliE, FlgB, FlgC, FlgF, FlgG). The basal body also contains a type III secretion system consisting of a transmembrane export gate complex (FlhA, FlhB, FliP, FliQ, FliR) and a cytoplasmic ATPase ring complex (FliH, FliI, FliJ). The MS-C-ring complex acts as a rotor of the flagellar motor. The P and L rings together form a very strong and stable ring complex and acts as a molecular bushing for high-speed rotation of the rod acting as a drive shaft. The C-ring, MS-ring, and LP-ring have 34-fold, 34-fold, and 26-fold rotational symmetries, respectively, whereas the rod is a helical assembly consisting of 11 protofilaments. The rod is divided into two structural parts: the proximal rod formed by 6 FliE subunits, 5 FlgB subunits, 6 FlgC subunits, and 5 FlgF subunits in this order; and the distal rod formed by 24 FlgG subunits. The proximal rod is firmly attached to the MS-ring and the polypeptide channel complex formed by 5 FliP subunits, 4 FliQ subunits, and 1 FliR subunit through interactions of FliE with FliF, FliP and FliR. OM, outer membrane; PG, peptidoglycan layer; CM, cytoplasmic membrane.

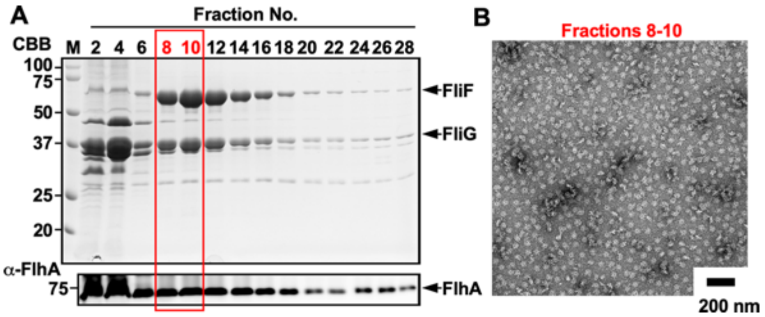

**FIG S2. Co-purification of the MS-ring with FliG and FlhA by sucrose gradient ultracentrifugation. (A)** SDS-PAGE band pattern of each fraction after sucrose gradient ultracentrifugation. After SDS-PAGE, proteins in each fraction were analyzed by Coomassie Brilliant blue staining and immunoblotting with polyclonal anti-FlhA antibody (α-FlhA). The positions of molecular mass markers (kDa) are shown on the left. **(B)** Negative stained EM image of purified MS-rings. Fractions 8, 9, and 10 containing FliF, FliG, and FlhA were collected and concentrated by ultracentrifugation. Concentrated samples were negatively stained with 2% (w/v) uranyl acetate and observed under an electron microscope.

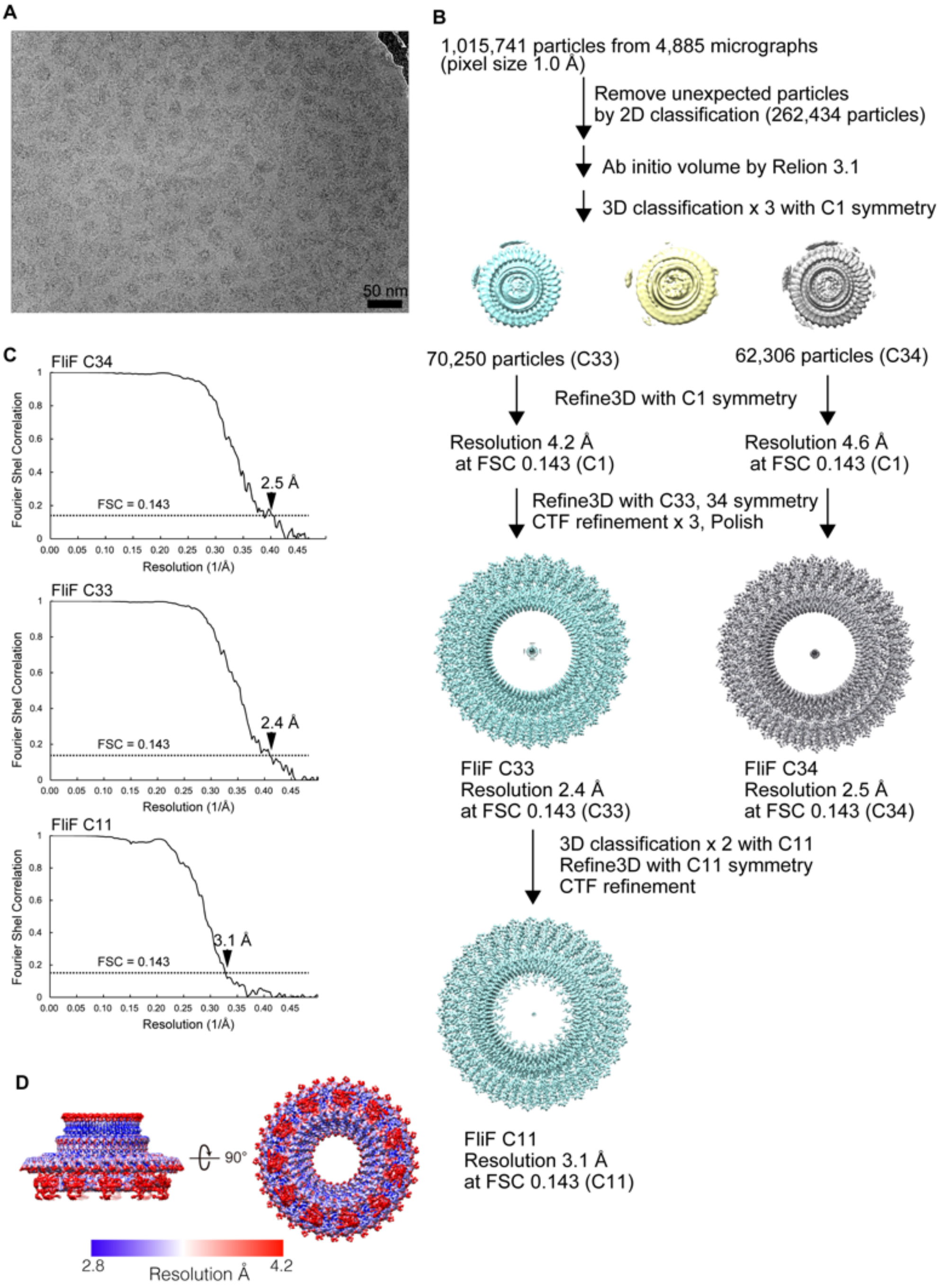

**FIG S3. CryoEM single particle 3D image analysis of the MS-ring. (A)** Representative cryoEM image of the MS-rings formed by co-expression of FliF with FliG and transmembrane export gate proteins. **(B)** Work process of cryoEM single particle 3D reconstruction. **(C)** Fourier shell correlation (FSC) curve of the final density map of the FliF-ring reconstructed with C34 (EMDB ID: EMD-60009) (upper), C33 (EMDB ID: EMD-60008) (middle), or C11 (EMDB ID: EMD-60007) (lower) symmetry applied. **(D)** Local resolution of the final density map of the FliF-ring with C11 symmetry applied is colored from blue (2.8 Å) to red (4.2 Å).

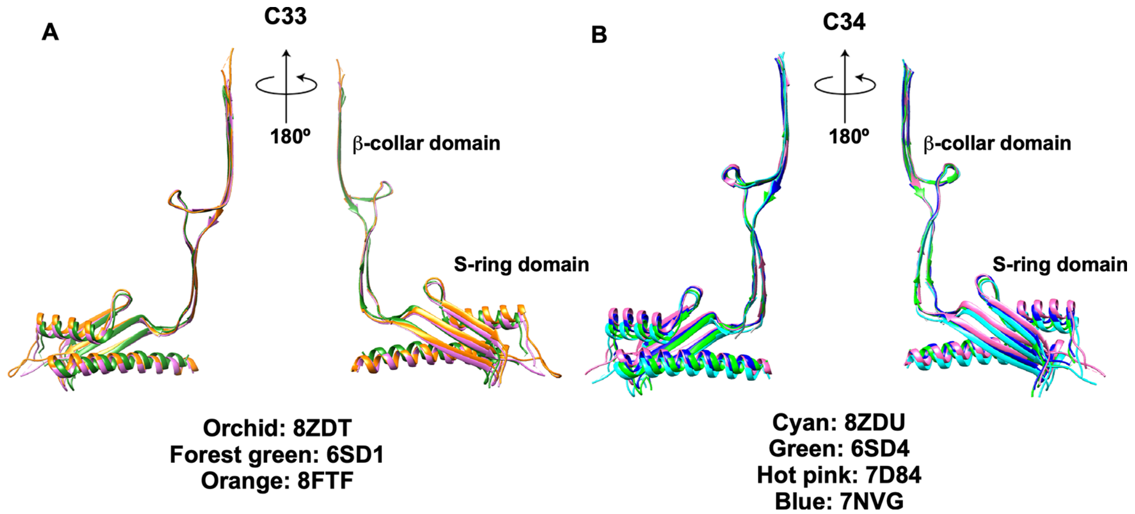

**FIG S4. Structural comparison of various *Salmonella* FliF subunits. (A)** The 33-mer RBM3-ring. The β-collar domain of the FliF subunit of the 6SD1 (forest green) and 8FTF (orange) ring structures were superimposed on to the corresponding domain of the coordinates obtained in this study (PDB ID: 8ZDT). **(B)** The 34-mer RBM3-ring. The β-collar domain of the FliF subunit of the 6SD4 (green), 7D84 (hot pink), and 7NVG (blue) ring structures were superimposed on to the equivalent domain of the coordinates obtained in this study (PDB ID: 8ZDU).

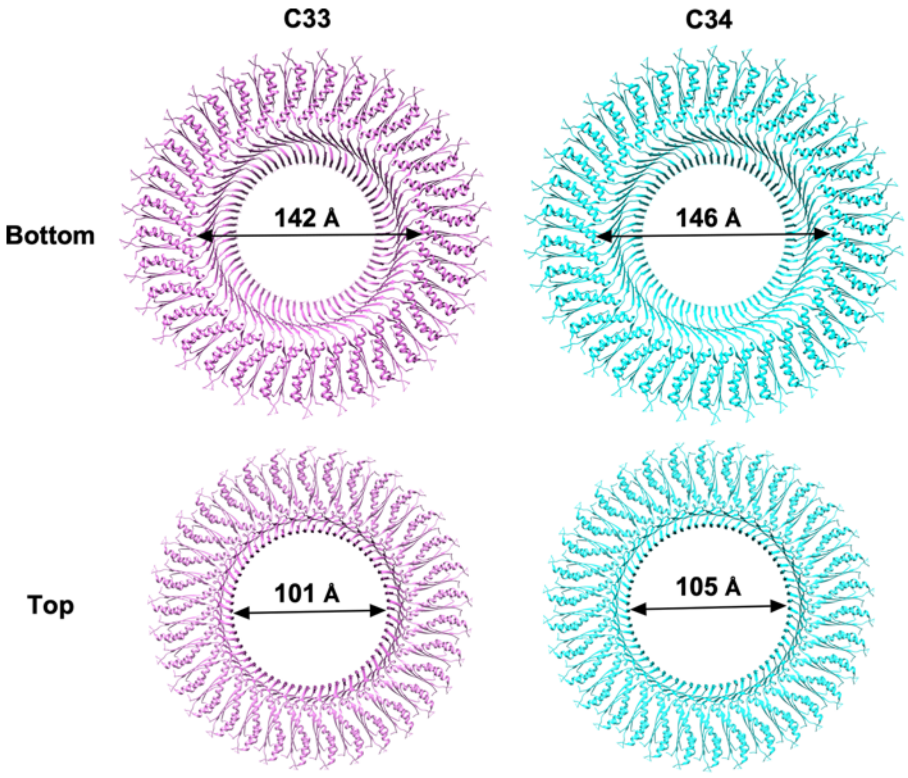

**FIG S5. Inner diameters of the 33-mer (PDB ID: 8ZDT) (left panels) and C34-mer (PDB ID: 8ZDU) (right panels) RBM3-rings.** The distance between Asp-229 of Mol-A and Asp-229 of Mol-Q and that between Asp-229 of Mol-A and Asp-229 of Mol-R were measured as the inner diameter of the 33-mer and 34-mer S-ring, respectively (upper panels). The distance between Pro-355 of Mol-A and Pro-355 of Mol-Q and that between Pro-355 of Mol-A and Pro-355 of Mol-R were measured as the inner diameter of the 33-mer and 34-mer β-collar, respectively (lower panels).

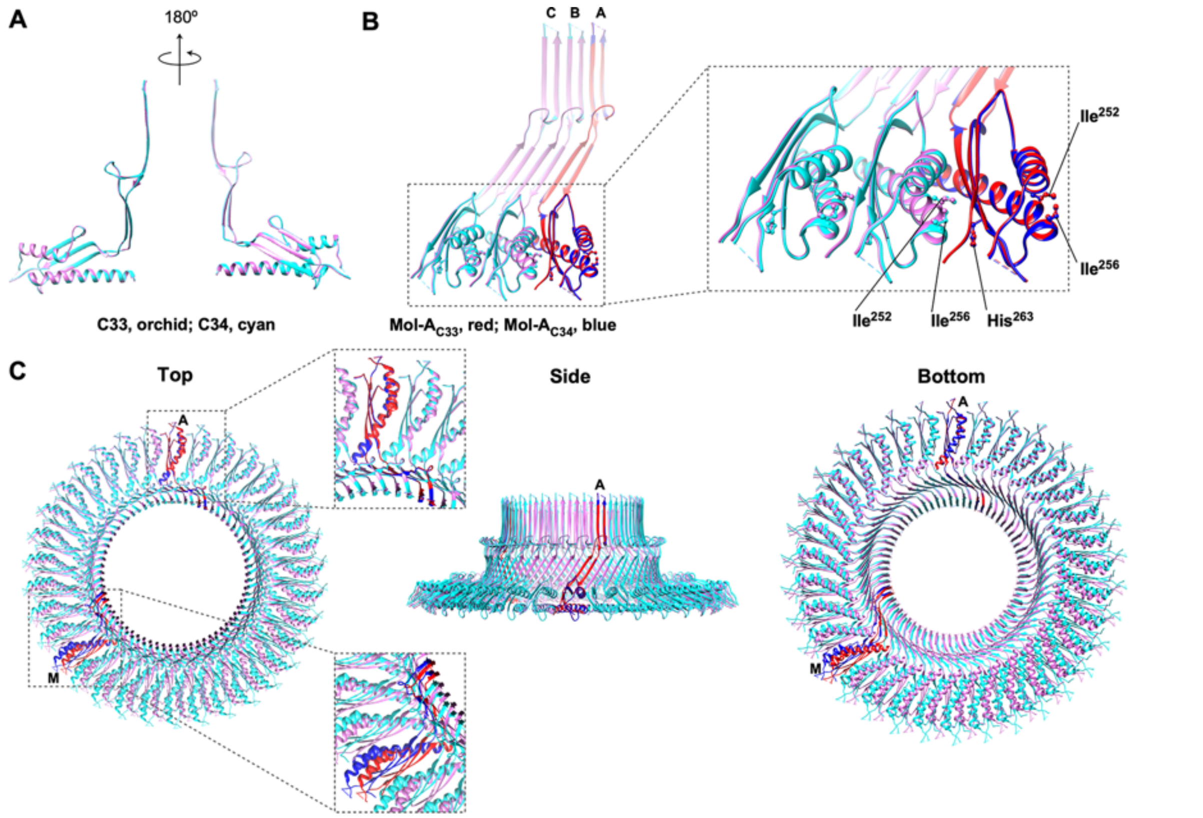

**FIG S6. Structural comparison of the 33-mer and 34-mer RBM3-rings. (A)** Structural comparison of Mol-A subunits in the 33-mer (PDB ID: 8ZDT) (orchid) and 34-mer (PDB ID: 8ZDU) (cyan) rings. Mol-A (blue) of the 34-mer ring was superimposed on Mol-A (red) of the 33-mer ring. **(B)** Intermolecular interface between FliF subunits. Only three FliF subunits (Mol-A, Mol-B, Mol-C) are shown. Residues Ile-252, Ile-256, and His-263 are located at the interface between subunits in the S-ring. **(C)** Superposition of the 33-mer and 34-mer rings.

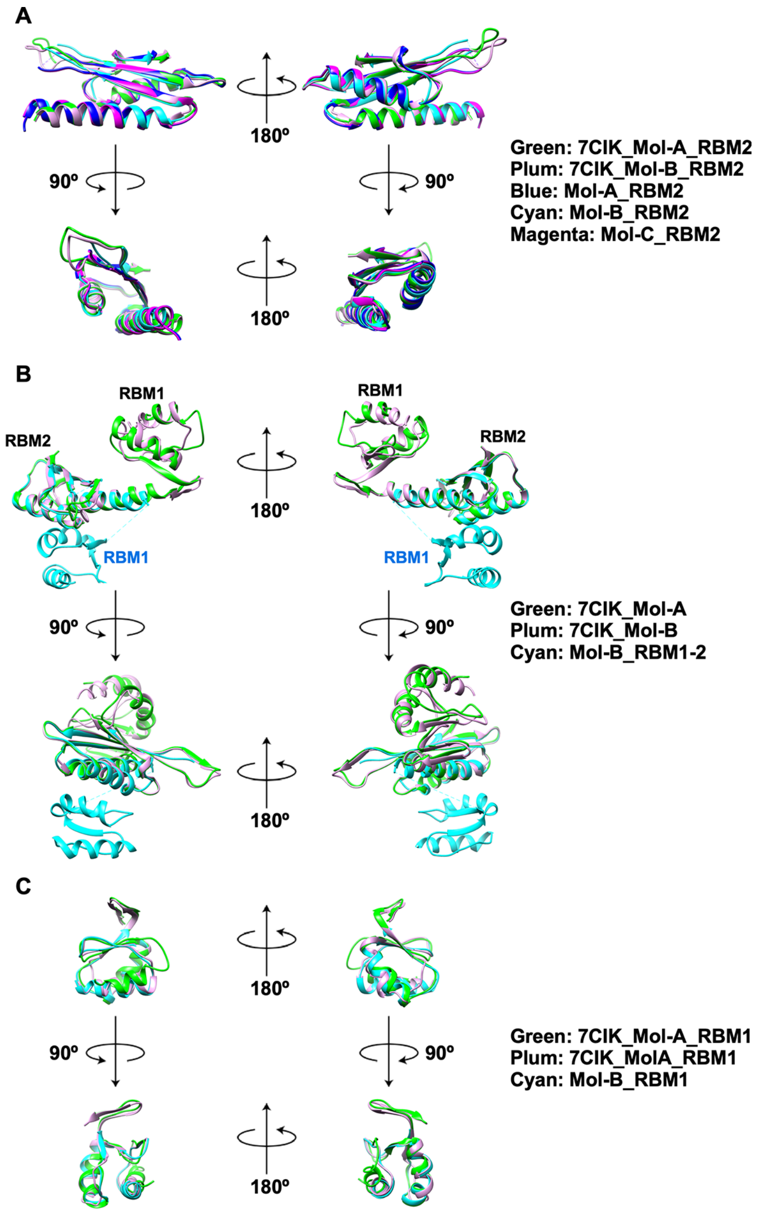

**FIG S7 Structural comparison of each domain of *Salmonella* FliF in the ring structure (PDB ID: 8ZDS) with those of *Aquifex* FliF crystal structure (PDB ID: 7CIK), which contains two molecules in an asymmetric unit (Mol-A, green; Mol-B, plum). (A)** Structural comparison of RBM2 in the inner core ring and RBM2 of cog-like structure of *Salmonella* FliF with the two RBM2 domains of *Aquifex* FliF. They are all nearly identical to one another. **(B)** Structural comparison of RBM1-RBM2. The position of RBM1 relative to RBM2 in *Salmonella* FliF is very different from those in *Aquifex* FliF. **(C)** Structural comparison of RBM1. They are almost identical to one another although slight differences are observed.

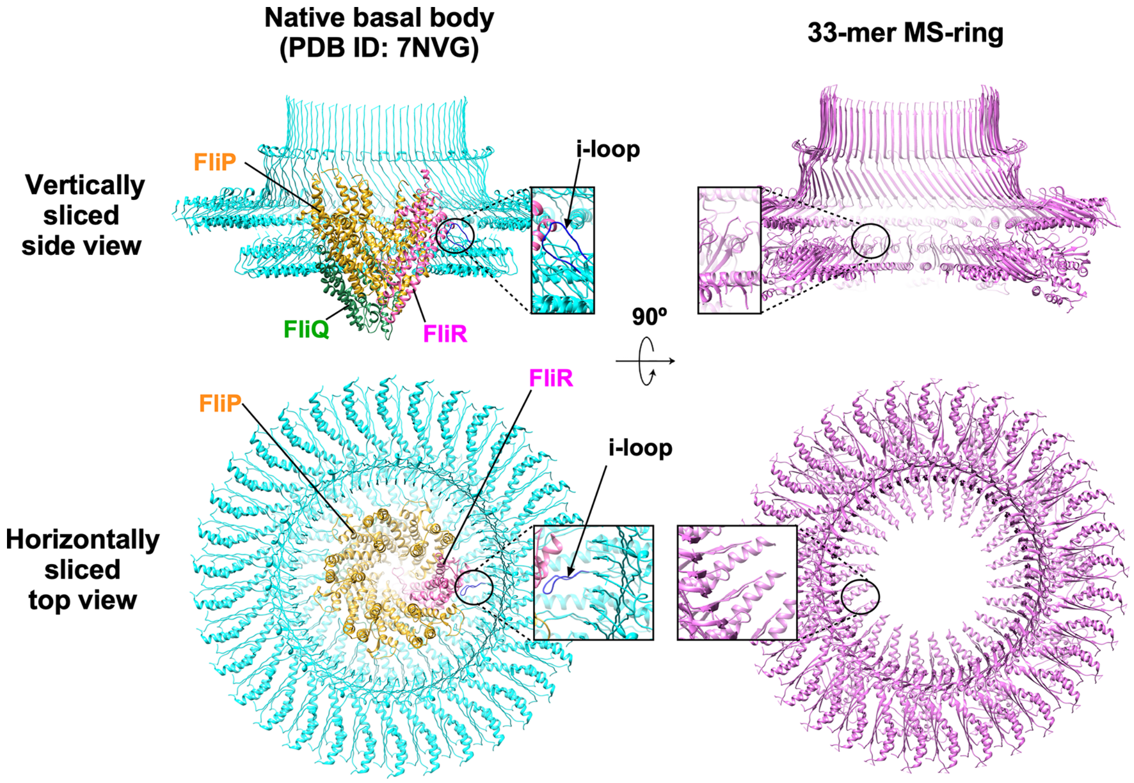

**FIG S8. Comparison of the i-loop conformations in the native basal body MS-ring (PDB ID: 7NVG) (left) and our 33-mer MS-ring with C11 symmetry applied (PDB ID: 8ZDS) (right).** Five FliP subunits (goldenrod) and one FliR subunit (hot pink) assemble into the $FliP_5$-$FliR_1$ complex, and then four FliQ subunit (sea green) surround the $FliP_5$-$FliR_1$ complex. The $FliP_5$-$FliQ_4$-$FliR_1$ complex is formed with a helical array of subunits and is located in the central pore of the RBM2-ring in the native basal body MS-ring. The i-loop of each RBM2 domain (residues 159–172) associates with either FliP or FliR subunit, and hence residues 163–168 are visible (left panels). However, these residues are invisible in our structure. Furthermore, the RBM1 domains below the inner core RBM2-ring are missing in both MS-ring structures.

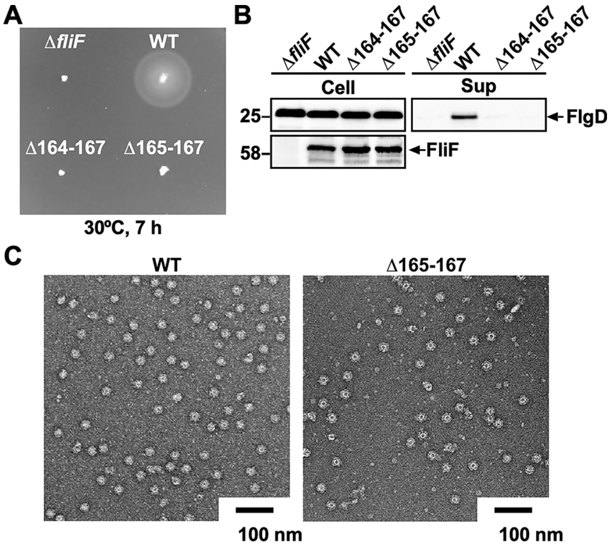

**FIG S9. Effect of small in-frame deletions within the i-loop of RBM2 on flagellar assembly. (A)** Motility of a *Salmonella fliF* null mutant harboring pTrc99AFF4 (Δ*fliF*), pMKMiF015 (WT), pMKMiF015(Δ164–167) (indicated as Δ164–167), or pMKMiF015(Δ165–167) (indicated as Δ165–167) in soft agar. The plate was incubated at 30ºC for 7 hours. **(B)** Secretion assays. Whole cell proteins (Cell) and culture supernatant fractions (Sup) were prepared from the above transformants. A 5 μl solution of each protein sample, which was normalized to an optical density of $OD_{600}$, was subjected to SDS-PAGE, followed by immunoblotting with polyclonal anti-FlgD (first row) or anti-FliF (second row) antibody. The positions of molecular mass markers (kDa) are shown on the left. **(C)** Negative stained EM images of the MS-rings isolated from the WT and Δ165–167 cells.

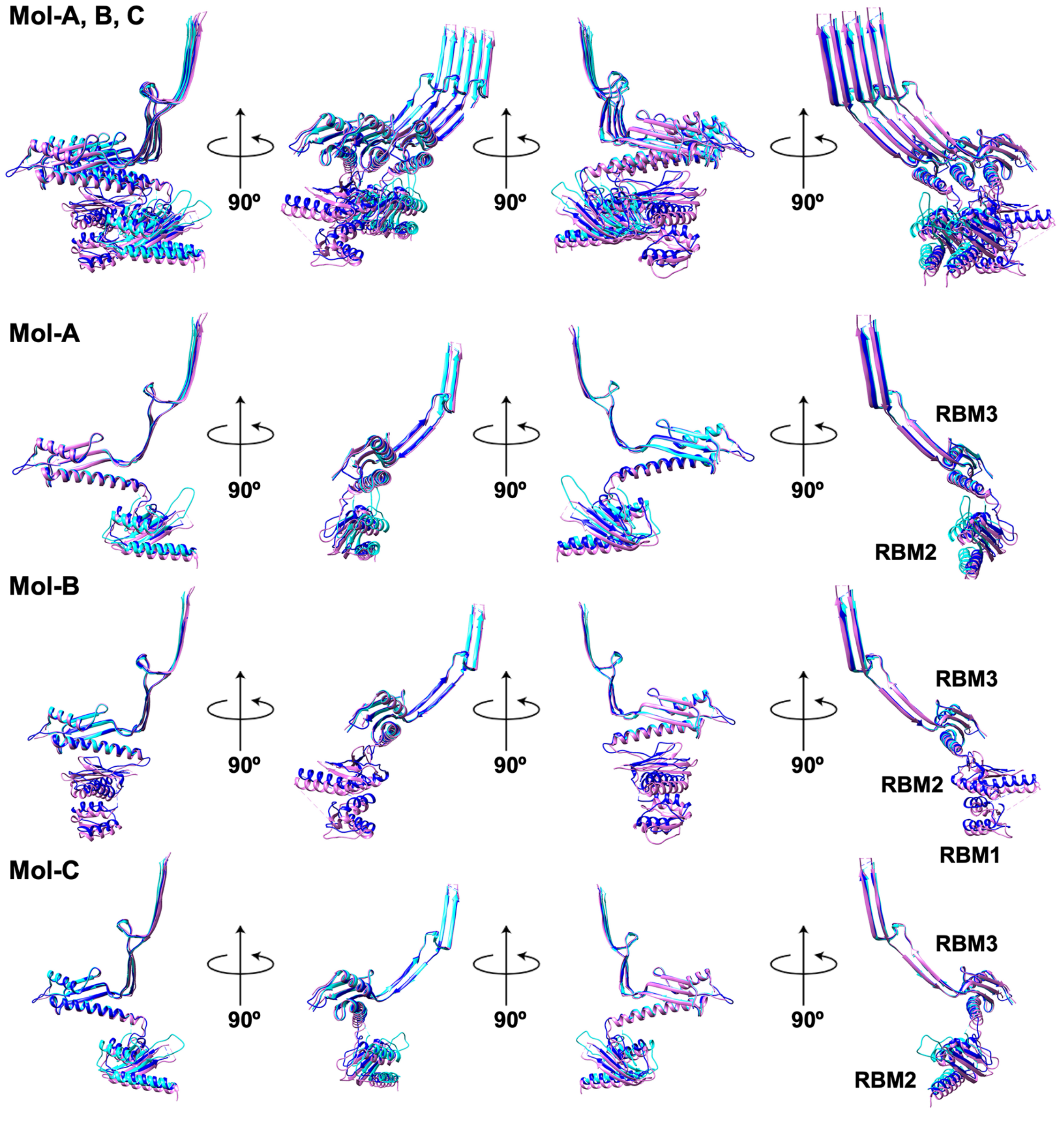

**FIG S10. Superposition of three FliF subunits obtained in this study (PDB ID: 8ZDS) (orchid) with equivalent coordinate regions of the 8T8P (blue) and 7NVG (cyan) structures.** The β-collar domain of one of the FliF subunits of the 8T8P (blue) or 7NVG (cyan) ring was superimposed on to the corresponding domain of Mol-A of the MS-ring obtained in this study to compare the structure of three subunits together as well as the conformation of each subunit.

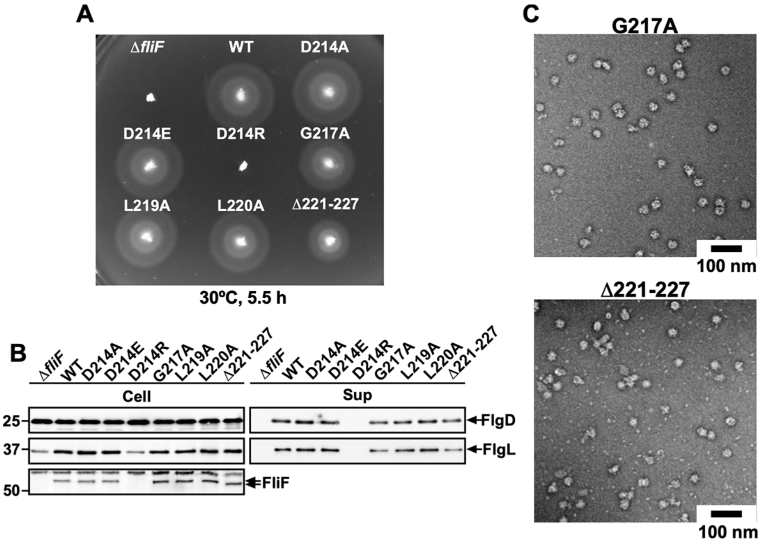

**FIG S11. Mutational analysis of the RBM2-3 loop connecting RBM2 and RBM3. (A)** Motility of a *Salmonella fliF* null mutant harboring pTrc99AFF4 (indicated as △*fliF*), pMKMiF015 (indicated as WT), pMKMiF015(D214A) (indicated as D214A), pMKMiF015(D214E) (indicated as D214E), pMKMiF015(D214R) (indicated as D214R), pMKMiF015(G217A) (indicated as G217A), pMKMiF015(L219A) (indicated as L219A), pMKMiF015(L220A) (indicated as L220A), or pMKMiF015(∆221–227) (indicated as ∆221–227) in soft agar. The plate was incubated at 30ºC for 5.5 hours. **(B)** Secretion assays. Whole cell proteins (Cell) and culture supernatant fractions (Sup) were prepared from the above transformants. A 5 µl solution of each protein sample, which was normalized to an optical density of $OD_{600}$, was subjected to SDS-PAGE, followed by immunoblotting with polyclonal anti-FlgD (first row), anti-FlgL (second row) or anti-FliF (third row) antibody. The positions of molecular mass markers (kDa) are shown on the left. **(C)** Negative stained EM images of the MS-rings isolated from the G217A and ∆221–227 mutants.

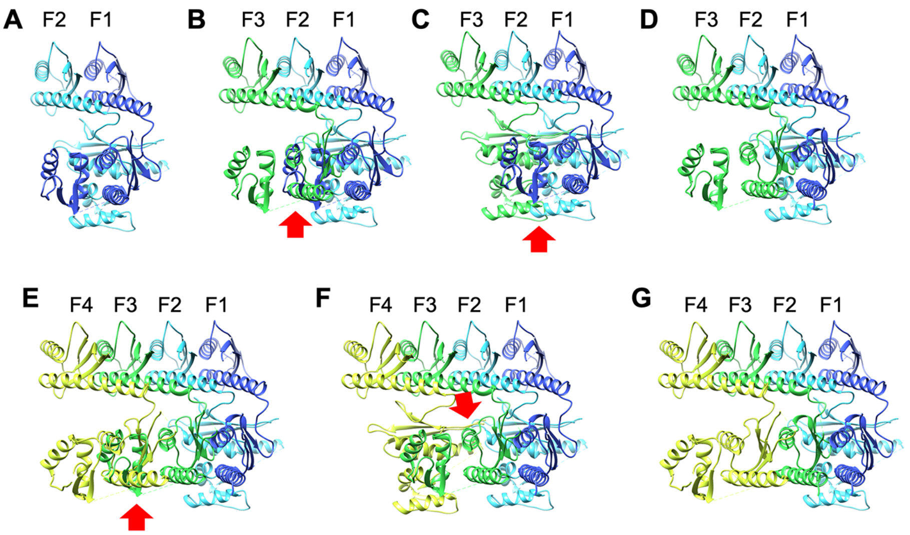

**FIG S12. Ribbon representation of the CCW growth model of MS-ring formation.** **(A)** Ribbon model of two RBM1-RBM2-RBM3 units (RBM1-3) of the initial FliF dimer (F1 in blue and F2 in cyan). **(B)** When the third subunit (F3 in green) is placed to the left of F2, serious steric hinderance occurs between RBM1 of F1 and RBM2 of F3 (shown by red arrow). **(C)** When RBM1-2 of F3 is placed in the conformation of Mol-B of the 33-mer MS-ring, steric hinderance also occurs between two RBM1-2 units of F2 and F3 (shown by red arrow). **(D)** Placement of RBM1-2 of F3 induces the dissociation of RBM1 from RBM2 in F1 to allow association of two RBM2 units of F1 and F3 to start forming the inner core RBM2-ring. **(E)** When the fourth subunit (F4 in yellow) is placed on the left side of F3, serious steric hinderance occurs between RBM1 of F3 and RBM2 of F4 (shown by red arrow). **(F)** When RBM1-2 of F4 is placed in the conformation of Mol-B, steric hinderance occurs again between two RBM1-2 units of F2 and F4 (shown by red arrow). **(G)** Placement of RBM1-2 of F4 forces RBM1 to detach from RBM2 in F3 to allow RMB2 of F4 to bind to RBM2 of F3 to grow the RBM2-ring.

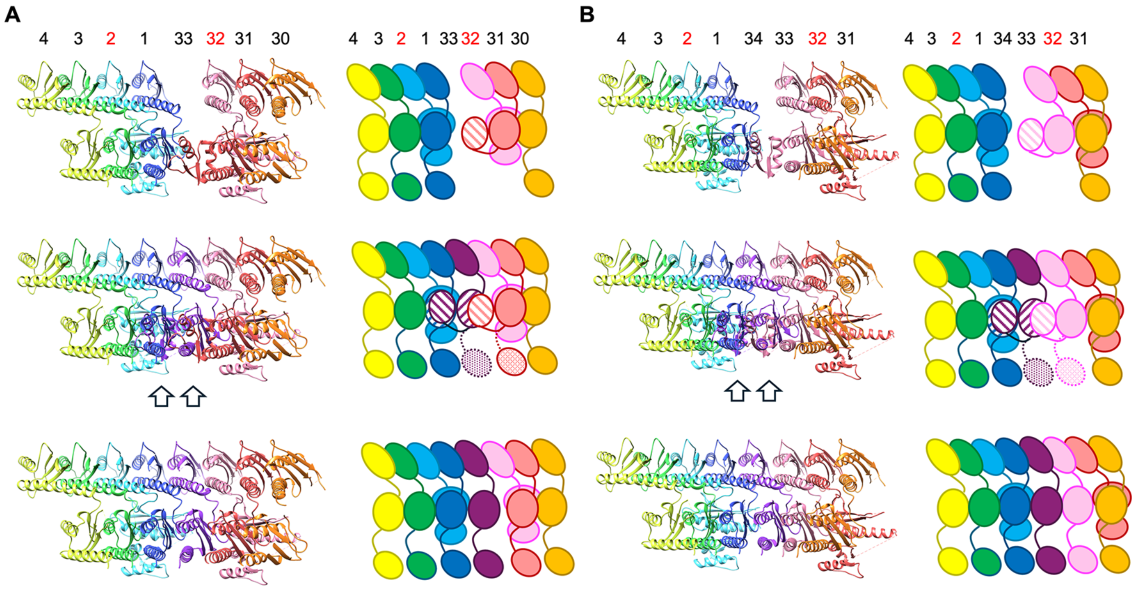

**FIG S13. Model for the final step of MS-ring formation. (A)** The final step of 33-mer MS-ring formation and **(B)** 34-mer MS-ring formation are shown in ribbon (left) and schematic representations (right). Top panels, before insertion of the final subunit; Middle panels, steric hindrance by the insertion of the final subunit; Bottom panels, completion of MS-ring formation. The subunit numbers are shown above each model.

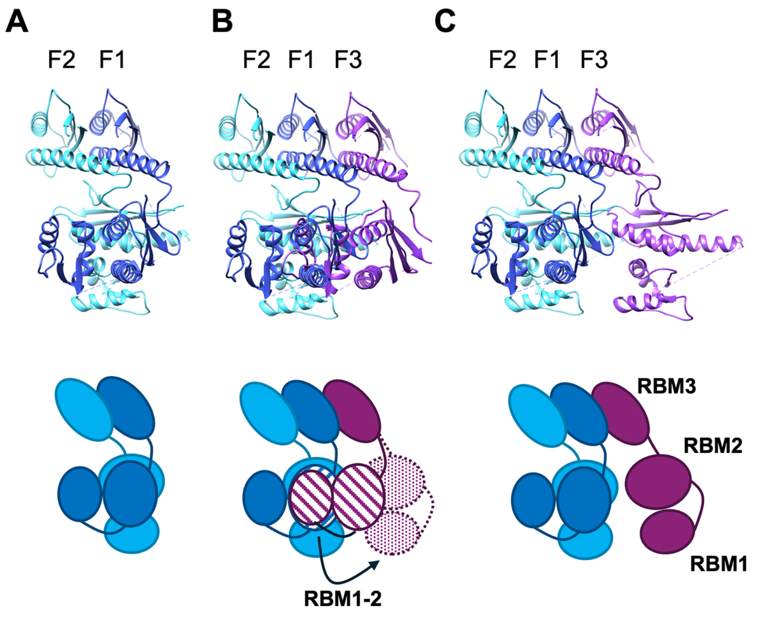

**FIG S14. CW growth model of MS-ring formation. (A-C)** Ribbon (top) and schematic (bottom) representations of the CW growth model. **(A)** Model for the RBM1-3 unit of the initial FliF dimer. **(B)** When the third subunit (F3) binds to the initial dimer on the right, steric hinderance occurs between RBM1 of F3 and RBM2 of F1 (shown by stripe pattern), and therefore, RBM1-2 of F3 move outward to avoid the clash but cannot be stabilized due to the lack of it binding site. **(C)** RBM1-2 of F3 moves freely, and no further ring growth occurs. Thus, MS-ring formation does not proceed in the CW direction.

**Table S1. EM data collection, processing, and refinement statistics.**

| | FliF C11 | FliF C33 | FliF C34 |
|---|---|---|---|
| EMDB | EMD-60007 | EMD-60008 | EMD-60009 |
| PDB | 8ZDS | 8ZDT | 8ZDU |
| **Data collection and processing** | | | |
| Magnification | 50,000 | 50,000 | 50,000 |
| Voltage (kV) | 300 | 300 | 300 |
| Electron exposure (e–/Å$^2$) | 40 | 40 | 40 |
| Defocus range (μm) | −0.5 – −2.5 | −0.5 – −2.5 | −1.0 – −2.5 |
| Pixel size (Å) | 1.00 | 1.00 | 1.00 |
| Symmetry imposed | C11 | C33 | C34 |
| Initial particle images (no.) | 757,555 | 757,555 | 757,555 |
| Final particle images (no.) | 68,598 | 216,718 | 194,233 |
| Map resolution (Å) | 3.1 | 2.5 | 2.6 |
| FSC threshold | 0.143 | 0.143 | 0.143 |
| **Refinement** | | | |
| Initial model used (PDB code) | 7D84, 7CIK | 7D84 | 7D84 |
| Model resolution | 3.70, 2.29 | 3.70 | 3.70 |
| FSC threshold | 0.143 | 0.143 | 0.143 |
| **Model composition** | | | |
| Non-hydrogen atoms | 69498 | 40623 | 42568 |
| Protein residues | 9053 | 5148 | 5304 |
| Ligands | 0 | 0 | 0 |
| ***B* factors (Å$^2$)** | | | |
| Protein | 163.48 | 38.82 | 44.73 |
| Ligand | 0 | 0 | 0 |
| **R.m.s. deviations** | | | |
| Bond length (Å) | 0.004 | 0.002 | 0.001 |
| Bond angles (°) | 0.703 | 0.444 | 0.446 |
| **Validation** | | | |
| MolProbity score | 1.73 | 1.12 | 1.16 |
| Clash score | 7.65 | 3.25 | 3.76 |
| Rotamer outliers(%) | 0.14 | 0.72 | 0.00 |
| **Ramachandran plot** | | | |
| Favored (%) | 95.46 | 99.33 | 99.33 |
| Allowed (%) | 4.54 | 0.67 | 0.67 |
| Disallowed (%) | 0 | 0 | 0 |

**Table S2. Inner diameters of the S-ring and β-collar**

| PDB ID | Inner diameter of the S-ring (Å) | Inner diameter of the β-collar (Å) |
|---|---|---|
| **C33** | | |
| 8ZDT (This study) | 141 | 101 |
| 6SD1 | 138 | 98 |
| 8FTF | 141 | 104 |
| **C34** | | |
| 8ZDU (This study) | 146 | 105 |
| 6SD4 | 140 | 102 |
| 7D84 | 144 | 103 |
| 7NVG | 142 | 103 |

**Table S3. Strains and plasmids used in this study**

| ***Salmonella* strains and Plasmids** | **Relevant characteristics** | **Source or reference** |
|---|---|---|
| *Salmonella* | | |
| SJW1368 | Δ(*cheW-flhD*); master operon mutant | 1 |
| TH12415 | Δ*fliF* | K. T. Hughes |
| Plasmids | | |
| pET3c | Expression vector | Novagen |
| pTrc99AFF4 | Expression vector | 2 |
| pMKM20001 | pTrc99CES3/ FliO + His-FliP + HA-FliQ + FliR-FLAG + FlhA + FlhB + FliF + FliG | 3 |
| pMKMiF001 | pET3c/ FliF | 4 |
| pMKMiF002 | pET3c/ FliF(I252A) | 4 |
| pMKMiF002-SP1 | pET3c/ FliF(I252A/R228C) | This study |
| pMKMiF002-SP2 | pET3c/ FliF(I252V) | This study |
| pMKMiF015 | pTrc99AFF4/ FliF | This study |
| pMKMiF015(Δ164-167) | pTrc99AFF4/ FliF(Δ164-167) | This study |
| pMKMiF015(Δ165-167) | pTrc99AFF4/ FliF(Δ165-167) | This study |
| pMKMiF015(Δ214-220) | pTrc99AFF4/ FliF(Δ214-220) | This study |
| pMKMiF015(Δ221-227) | pTrc99AFF4/ FliF(Δ221-227) | This study |
| pMKMiF015(D214A) | pTrc99AFF4/ FliF(D214A) | This study |
| pMKMiF015(D214E) | pTrc99AFF4/ FliF(D214E) | This study |
| pMKMiF015(D214R) | pTrc99AFF4/ FliF(D214R) | This study |
| pMKMiF015(G217A) | pTrc99AFF4/ FliF(G217A) | This study |
| pMKMiF015(G217P) | pTrc99AFF4/ FliF(G217P) | This study |
| pMKMiF015(G217W) | pTrc99AFF4/ FliF(G217W) | This study |
| pMKMiF015(L219A) | pTrc99AFF4/ FliF(L219A) | This study |
| pMKMiF015(L220A) | pTrc99AFF4/ FliF(L220A) | This study |
